# The Fc-effector function of COVID-19 convalescent plasma contributes to SARS-CoV-2 treatment efficacy in mice

**DOI:** 10.1101/2022.06.10.495677

**Authors:** Irfan Ullah, Guillaume Beaudoin-Bussières, Kelly Symmes, Marc Cloutier, Eric Ducas, Alexandra Tauzin, Annemarie Laumaea, Philippe Bégin, Walther Mothes, Priti Kumar, Renée Bazin, Andrés Finzi, Pradeep D. Uchil

## Abstract

COVID-19 convalescent plasmas (CCPs) are chosen for plasma therapy based on neutralizing titers and anti-Spike immunoglobulin levels. However, specific CCP characteristics that promote SARS-CoV-2 control in recipients are complex and incompletely defined. Using an *in vivo* imaging approach, we demonstrate that CCPs with low neutralizing and high Fc-effector activity, in contrast to those with poor Fc-function, afford effective prophylaxis and therapy in K18-hACE2 mice lethally challenged with SARS-CoV-2-nLuc. Macrophages and neutrophils significantly contributed to CCP effects during therapy but to a reduced extent under prophylaxis. Both IgG and Ig(M+A) were required during therapy, but the IgG fraction alone was sufficient during prophylaxis. Finally, despite neutralizing poorly, SARS-CoV-2 Wuhan-elicited CCPs delayed Delta and Beta variants of concern (VOC)-induced mortality in mice illustrating the contribution of polyclonal Fc-effector functions in immunity against VOCs. Thus, in addition to neutralization, Fc-effector activity is a significant criterion for CCP selection for therapeutic applications.

## Introduction

Convalescent plasma (CP) therapy is a first line of treatment when the human population lacks pathogen-specific immunity and treatment options are limited. CP therapy involves collecting plasma from recovered patients to harness “passive immunity” provided by pathogen-specific antibodies for preventing or treating disease in infected patients (Casadevall and Scharff, 1995). The use of CP therapy dates back to the 1918 influenza pandemic and more recently to fight outbreaks caused by SARS-CoV-1, MERS and Ebola (Cao and Shi, 2020; Luke et al., 2006; Mair-Jenkins et al., 2015). CP therapy may be of particular interest for the aged and immune-suppressed cancer or transplant patients where vaccination fails to elicit protective antibody responses as well as in co-morbid populations where vaccination cannot be used (Beraud et al., 2022; Ljungquist et al., 2022; Pinkus and Said, 1986). Unlike with vaccines and monoclonal antibodies (mAbs), CP therapy requires limited development and standard infrastructure for blood collection and is rapidly deployable globally even under low resource settings. CP is adaptable to emerging SARS-CoV-2 variants of concern (VOCs) when the plasma source is from convalescent human subjects infected with homologous variant virus strains. Additionally, the polyclonal nature of CPs also makes them relatively effective against heterologous variants. In contrast, targeted immune therapies need development from scratch to specifically target newly arising mutations as is currently the case with mRNA vaccines and neutralizing antibody (nAb) cocktails for SARS-CoV-2 Omicron variants and sublineages (Greaney et al., 2022; Tada et al., 2022; Tartof et al., 2022). Therefore, CP therapy remains a rapidly deployable go-to countermeasure for emerging and future pathogens with pandemic potential.

Currently, the choice of COVID-19 convalescent plasma (CCP) for therapy is driven by high titers of anti-SARS-CoV-2 Spike IgG (Median titer: 1:3200) and neutralization titer [inhibitory dilution (ID_50_] >1:250) (Villa, 2021). Neutralizing antibodies (nAbs) in CCPs can inactivate virus and CCP administration has reduced inflammation and helped mitigate SARS-CoV-2-induced acute respiratory disorder syndrome (ARDS) (Basheer et al., 2021). Thus, plasma neutralizing titers are a critical correlate of SARS-CoV-2 immunity and undoubtedly an important criterion for the selection of CCP for therapy (Dispinseri et al., 2021). However, the efficacy of CCP therapy has been questioned by large randomized clinical trials (RCTs) such as RECOVERY, CONCOR-1, and REMAP-CAP that found little to no evidence in reducing the risk of intubation or death from administration of high-titer CCP in hospitalized patients (Begin et al., 2021; Writing Committee for the et al., 2021). In the CONCOR-1 trial, higher levels of IgG specific for the membrane-bound Spike with disproportionally low neutralizing and Fc-effector functions were associated with worse outcomes (Begin et al., 2021). Emerging consensus from other RCTs is that if CCP is used, it should contain the highest neutralizing titers possible and be transfused early in the disease course before patients required greater supportive therapies to increase the likelihood of benefit (Korley et al., 2021).

CCPs are complex and contain an ensemble of polyclonal antibodies with different immunoglobulin (Ig) isotypes, epitope specificity, virus-neutralizing and non-neutralizing activities. Despite the presence of beneficial nAbs, the plasma milieu may or may not act in concert to produce the desired antiviral activities required for protection in recipients. Additional signatures of CCP that track with positive outcome are required to better characterize the clinical utility of and choice of CCP for plasma therapy. In addition to directly neutralizing, antibodies can utilize their Fc domain for mediating effector functions by interacting with Fc receptors (FcRs) expressed on innate immune cells (Beaudoin-Bussieres et al., 2022; Ullah et al., 2021; van Erp et al., 2019; Winkler et al., 2021). FcR engagement on neutrophils, monocytes, and natural killer (NK) cells can elicit multiple activities including the clearance of viral particles through phagocytosis (antibody-dependent phagocytosis; ADP) and cytotoxic killing of virus-infected cells (antibody-dependent cellular cytotoxicity; ADCC). However, not all antigen-binding nAbs or non-neutralizing antibodies (non-nAbs) have the ability to stimulate Fc effector functions. A multitude of factors including epitope specificity, angle of approach, isotype, glycosylation pattern, polymorphism and levels of FcR on innate immune cell type influence activation by antibody-antigen complexes (Lu et al., 2018; Patel et al., 2019; Pereira et al., 2018; Tay et al., 2019). Several studies have now shown that purified monoclonal SAR-CoV-2 nAbs rely on Fc-effector functions for improved *in vivo* efficacy especially during therapy (Schafer et al., 2021; Ullah et al., 2021; Winkler et al., 2021; Yamin et al., 2021). Moreover, introducing Fc-FcγR binding enhancer mutations (GASDALIE) have improved *in vivo* nAb efficacy and reduced dosage (Li et al., 2022; Yamin et al., 2021). 80-96% of Spike-binding antibodies in plasma are non-neutralizers (Jennewein et al., 2021). Given that the predominant proportion of antibodies in plasma are non-nAbs, their contribution to the overall Fc-mediated targeting of SARS-CoV-2 virions and virus-infected cells is expected to be significant. Non-nAbs, through Fc-function, may synergize with nAbs to improve overall efficacies especially in a polyclonal setting like in the plasma (Beaudoin-Bussieres et al., 2022; Tauzin et al., 2021). Furthermore, Fc-functions of antibodies elicited by prior infection or vaccination are a suggested correlate for continued immunity against emerging variants of concerns (VOCs) despite compromised neutralization (Anand et al., 2021; Kaplonek et al., 2022b; Richardson et al., 2022; Tauzin et al., 2021). Thus, given the emerging evidence of Fc-mediated antibody effector functions in both protection and disease caused by SARS-CoV-2, presence of robust Fc-effector activities may serve as an additional criterion for selecting CCPs for therapeutic applications. While the presence of SARS-CoV-2-specific antibodies in CCPs that elicit Fc-mediated effector activity have independently correlated with therapeutic benefits (Begin et al., 2021), direct *in vivo* evidence beyond correlation is lacking.

K18-hACE2 mice are highly susceptible to SARS-CoV-2 infection (Halfmann et al., 2022; McCray et al., 2007; Park et al., 2022; Seehusen et al., 2022; Zheng et al., 2021). They represent a practical and relevant animal model to rapidly navigate through the multiple complex activities of CCP and identify those that contribute to protection, addressing limitations of *in vitro* assay-driven plasma analyses that cannot predict *in vivo* effects. Here we used the K18-hACE2 mouse model together with bioluminescence imaging for tracking SARS-CoV-2 infection to screen CCPs. Our screen identified a CCP with low neutralizing (ID_50_<1:250) but robust Fc-effector activity that protected mice from lethal challenge with homologous WA1 strain prophylactically as well as therapeutically. In contrast, CCPs with similarly low neutralizing titers but poor Fc-effector activity did not confer protection. Depletion of macrophages and neutrophils revealed that these innate immune cells contributed significantly to CCP-mediated protection but to a higher extent during therapy than under prophylaxis. Depletion of antibody classes from plasma showed that IgG as well as Ig(M+A) fractions were required for maximal *in vivo* efficacy during therapy. In contrast, the IgG fraction alone sufficed for prophylaxis. IgG-Fc-effector functions were however crucial for prophylaxis in the absence of Ig(M+A). Furthermore, ancestral SARS-CoV-2 (Wuhan)-elicited CCPs delayed mortality by Delta and Beta VOCs despite sub-optimal neutralization demonstrating the importance of polyclonal Fc-effector functions in cross-immunity against VOCs. These data make a compelling case for the relevance of Fc-effector activities when assessing CCP therapeutic potency and suggest that it could potentially serve as an additional criterion for its selection. Our study, in addition to highlighting the versatility of the K18-mouse model in dissecting complex activity mechanisms of CCPs, identifies polyclonal Fc-effector functions as a key CCP profile necessary for the successful treatment of infections by SARS-CoV-2 and VOCs.

## Results

### Screening in SARS-CoV-2-challenged K18-hACE2 mice allows identification of COVID-19 convalescent plasmas (CCPs) with net protective profiles

We classified CCPs with low nAb titer [inhibitory dilution (ID_50_) ≤ 250] (Gundlapalli et al., 2021; Villa, 2021) collected during the first wave of COVID-19 based on their *in vitro* ADCC activity **(Figure 1A, B)**. To determine whether Fc activity translates to a protective profile *in vivo*, we analyzed CCPs for their effects in K18-hACE2 mice challenged with homologous SARS-CoV-2 WA1 under both prophylactic and therapeutic regimens **(Figure 1C)**. Analyses of body weight loss, N mRNA copy numbers and survival revealed that CCP-6, with the highest ADCC activity *in vitro* (%ADCC= 91), prevented body weight loss, controlled virus replication in target organs (nose, lungs and brain) and averted from SARS-CoV-2-induced mortality **(Figure 1D-F)**. Mice treated prophylactically with CCP-3, 4 and 5 with reasonable ADCC activity (%ADCC= 50-60%), lost body weight initially but started recovering by 7 dpi and eventually controlled virus replication in target organs. In contrast, mice that received CCP-1 or CCP-2 with low ADCC activity (11 and 16% respectively) did not control virus replication (N mRNA copy numbers) in target organs and succumbed by 6 dpi like the mouse in mock-treated cohort. The importance of Fc function was evident when 3 out of 4 CCPs with ADCC activity also rescued mice from SARS-CoV-2-induced mortality upon administration at 2 dpi after infection was established **(Figure 1G-I)**. To note however, CCP-5, that had the lowest neutralizing activity, did not protect therapeutically despite reasonable Fc activity. Accordingly, polyclonal Fc-effector functions may contribute to protection, but some level of neutralizing activity may be necessary for therapeutic control of established infections. Thus, our data highlights the complexity of plasma milieu where a single characteristic may not be sufficient to define candidate CCPs for therapy and showcases the utility of *in vivo* screening to identify CCPs with net protective characteristics for an optimal therapeutic outcome.

**Figure 1.**
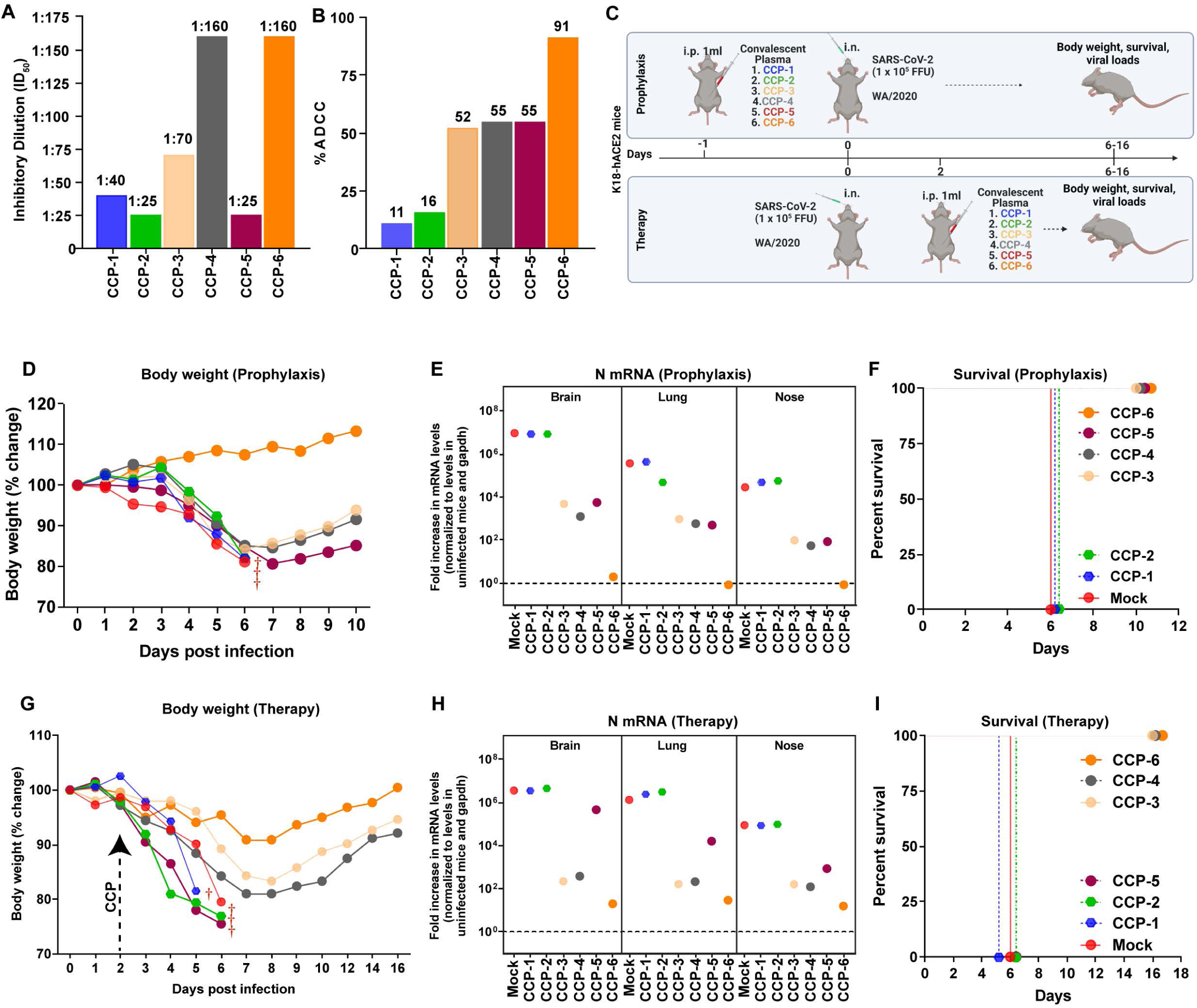
*In vivo* screening of COVID-19 Convalescent Plasma (CCPs) (A) A graph depicting WA1-neutralizing activity of indicated CCPs expressed as inhibitory dilution of plasma (ID_50_) that reduces FFUs by 50% using Vero E6 cells as targets. (B) %ADCC in the presence of CCP using a 1:1 ratio of parental CEM.NKr cells and CEM.NKr.Spike cells as target cells while PBMCs from uninfected donors were used as effector cells. (C) A scheme showing experimental design for screening *in vivo* efficacy of indicated CCPs delivered 1ml per 20-25 g body weight of mouse intraperitoneally (i.p.) under prophylaxis (-1dpi) and therapeutically (+2 dpi) in K18-hACE2 mice intranasally (i.n.) challenged with 1 x 10^5^ FFU WA1 SARS-CoV-2-nLuc. PBS-treated mice were used as control (Mock). (D, G) Temporal changes in mouse body weight with initial body weight set to 100% during CCP prophylaxis (-1dpi) and therapy (+2 dpi) for experiment as in C. (E, H) Fold change in SARS-CoV-2 nucleocapsid (N gene) expression in brain, lung and nose tissues during CCP prophylaxis and therapy for experiment shown in C. The data were normalized to *Gapdh* mRNA expression in the same sample and that in non-infected mice after necropsy. (F, I) Kaplan-Meier survival curves for evaluating *in vivo* efficacy of CCPs against SARS-CoV-2- nLuc for an experiment as in C.

### In-depth Analysis of Protective Effects of CCPs With Fc-effector Activity

To better understand the basis for CCP-mediated protection we chose CCP-6 whose protective profile depended on both neutralizing and Fc-effector functions as well as CCP-2 with low ADCC activity (16% vs 91% for CCP-6) as well as low neutralizing activity (1:25 vs 1:160 for CCP-6) for in-depth characterization and bioluminescence imaging (BLI)-based studies **(Figure 2)**. We first prophylactically treated mice (n=7) with CCPs (i.p.) 1 day before intranasal (i.n.) challenge with SARS-CoV-2-nLuc **(Figure 2A-F)**. Temporal BLI imaging and quantification of nLuc signals to monitor virus replication revealed that prophylaxis with CCP-2 or hIgG1 did not prevent SARS-CoV-2 WA1 nLuc infection and subsequent virus spread **(Figure 2B-D)**. In contrast, CCP-6 prophylaxis completely blocked initiation of virus infection as no detectable nLuc signals were observed in K18-hCE2 mice. These data were corroborated by body weight and survival analyses where isotype-treated control and CCP-2 treated mice steadily lost body weight upon infection and succumbed to infection by 6 dpi while CCP-6-treated mice gained weight indicating complete protection **(Figure 2E-F)**. nLuc signals measured separately in individual isolated target organs after necropsy also corresponded to viral loads (N mRNA expression, viral titers) **(Figure S1A-D)**. In addition, CCP-2 and isotype-treated control cohorts displayed 10-1000-fold induction of inflammatory cytokine mRNA expression in target organs **(Figure S1E-F)**. In contrast, cytokine mRNA expression in CCP-6 treated animals were at basal levels indicating complete protection from SARS-CoV-2 infection.

**Figure 2.**
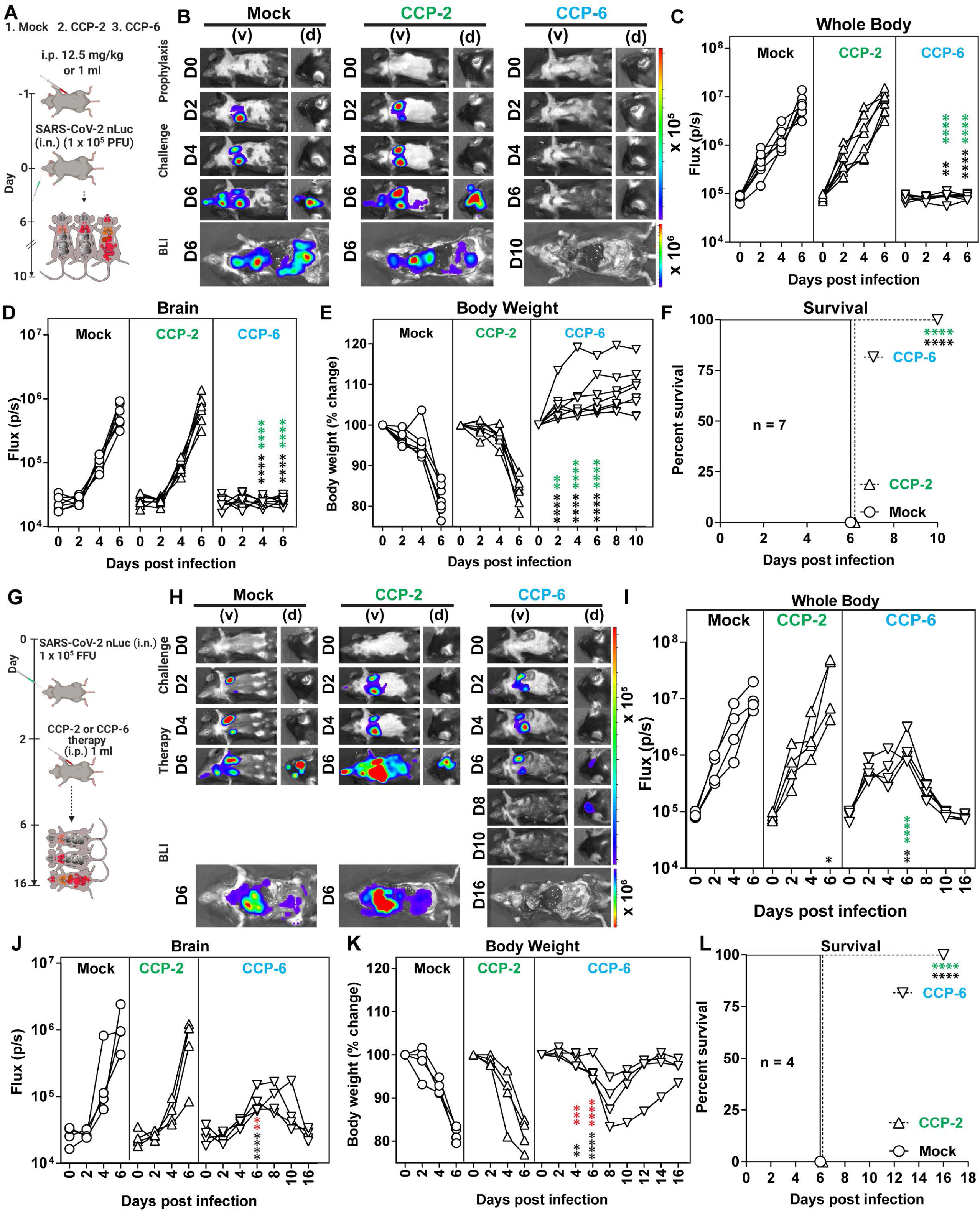
CCP with Fc-effector Activity Protects K18-hACE2 Mice Against Lethal SARS-CoV-2 Challenge During Prophylaxis and Therapy. (A, G) A scheme showing experimental design for testing *in vivo* efficacy of CCP-2 and CCP-6 (1 ml/ 20-25 g body weight, i.p.) under prophylaxis (-1 dpi) and therapy (+2 dpi) in K18-hACE2 mice intranasally challenged with 1 x 10^5^ FFU SARS-CoV-2-nLuc. hIgG1 isotype-treated mice were used as control (Mock). (B, H) Representative BLI images of SARS-CoV-2-nLuc-infected mice in ventral (v) and dorsal (d) positions for experiment as in A and G. Scale bars denote radiance (photons/sec/cm^2^/steradian). (C, D, I, J) Temporal quantification of nLuc signal as flux (photons/sec) computed non-invasively in indicated tissues. (E, K) Temporal changes in mouse body weight with initial body weight set to 100% for experiments shown in A and G. (F, L) Kaplan-Meier survival curves of mice (n = 7 and 4 per group) statistically compared by log-rank (Mantel-Cox) test for experiments as in A and G. Each curve in C-E and I-K represents an individual mouse. Data in panels C-F and I-L are from two independent experiments and n=2-4 mice per group. Grouped data in (C-E) and (I-K) were analyzed by 2-way ANOVA followed by Tukey’s multiple comparison tests. Statistical significance for group comparisons to mock controls are shown in black, with convalescent plasma CCP-2 are shown in green, and with CCP-6 are shown in cyan. ∗, p < 0.05; ∗∗, p < 0.01; ∗∗∗, p < 0.001; ∗∗∗∗, p < 0.0001; Mean values ± SD are depicted. See also Fig S1

To confirm the ability of CCP-6 to clear established infection (therapeutic mode), we treated mice 2 days after intranasal (i. n.) challenge with SARS-CoV-2-nLuc **(Figure 2G)**. Quantification of nLuc signals after temporal BLI imaging revealed that therapy with the CCP-2 or hIgG1 did not control expanding SARS-CoV-2 replication in the lungs and allowed virus dissemination into the brain in K18-hACE2 mice **(Figure 2H-J)**. In contrast, mice treated with CCP-6 cleared pre-established infection in the lungs by 8 dpi. Remarkably, despite detectable neuroinvasion at 6 dpi, CCP-6 treatment controlled and subsequently cleared virus in the brain of infected animals by 10 dpi **(Figure 2H-J)**. These data were again corroborated in body weight analyses and survival experiments where CCP-2 and hIgG1-treated mice lost ∼20% of their starting body weight and succumbed to infection by 6 dpi while all CCP-6-treated mice survived and regained body weight **(Figure 2K-L)**. A significant decrease in nLuc signal intensity was also seen in individual target tissues (nose, lung brain) post-necropsy in CCP-6 treated mice, which correlated with viral loads as well as mRNA levels of Nucleocapsid (N) and inflammatory cytokines in contrast to hIgG1 and CCP-2-treated cohorts of mice **(Figure S1G-L)**. Altogether, these data confirmed the importance CCP-6 with robust ADCC activity in conferring protection particularly against established SARS-CoV-2 infection, when CCPs are mainly deployed for therapeutic benefits.

### Macrophages and Neutrophils Contribute Marginally to Protection Mediated by CCP in Prophylaxis

CCP potency against SARS-CoV-2 is a result of both neutralizing activity and Fc-mediated mobilization of innate immune cells by antibodies for elimination of virus particles and infected cells. To evaluate involvement of innate immune cells in CCP-6 prophylaxis, we immuno-depleted neutrophils (anti-Ly6G) or macrophages (anti-CSF1R) during CCP-prophylaxis. Flow cytometry confirmed that ∼98% neutrophils (CD45^+^CD11b^+^Ly6G^+^) in blood or ∼75% of lung-resident macrophages (CD45^+^CD11b^+^ Ly6G^-^Ly6C^-^CD68^+^) were depleted following treatment with depleting antibodies **(Figure S2A-D)**. Depletion of these innate immune cell types did not alter the susceptibility of K18-hACE2 mice to SARS-CoV-2 infection **(Figure S2E-J)**. BLI analyses revealed transient and weak SARS-CoV-2 replication in the lungs at 4 and 6 dpi that cleared by 10 dpi in CCP-6-treated cohort with immune cell depletion, as determined by the nLuc signal (Flux p/s) **(Figure 3A-C)**. In addition, CCP-6 could still prevent virus dissemination to the brain **(Figure 3D)**. However, a transient body weight loss (up to 10%) in K18-hACE2 mice occurred before complete recovery in contrast to mice that were not depleted of these innate cell types **(Figure 3E, F)**. Post-necropsy analyses (organ flux, tissue viral loads and inflammatory cytokine mRNA expression) also confirmed virological control at the experimental endpoint **(Figure 3G-L)**. However, inflammatory cytokines were significantly higher in the lungs compared to undepleted cohorts. Thus, while the neutralizing activity played a major role during CCP-6 prophylaxis, our data indicate a marginal but distinct contribution of antibody Fc-effector functions in engaging immune cells to control residual infection and mitigating inflammation.

**Figure 3.**
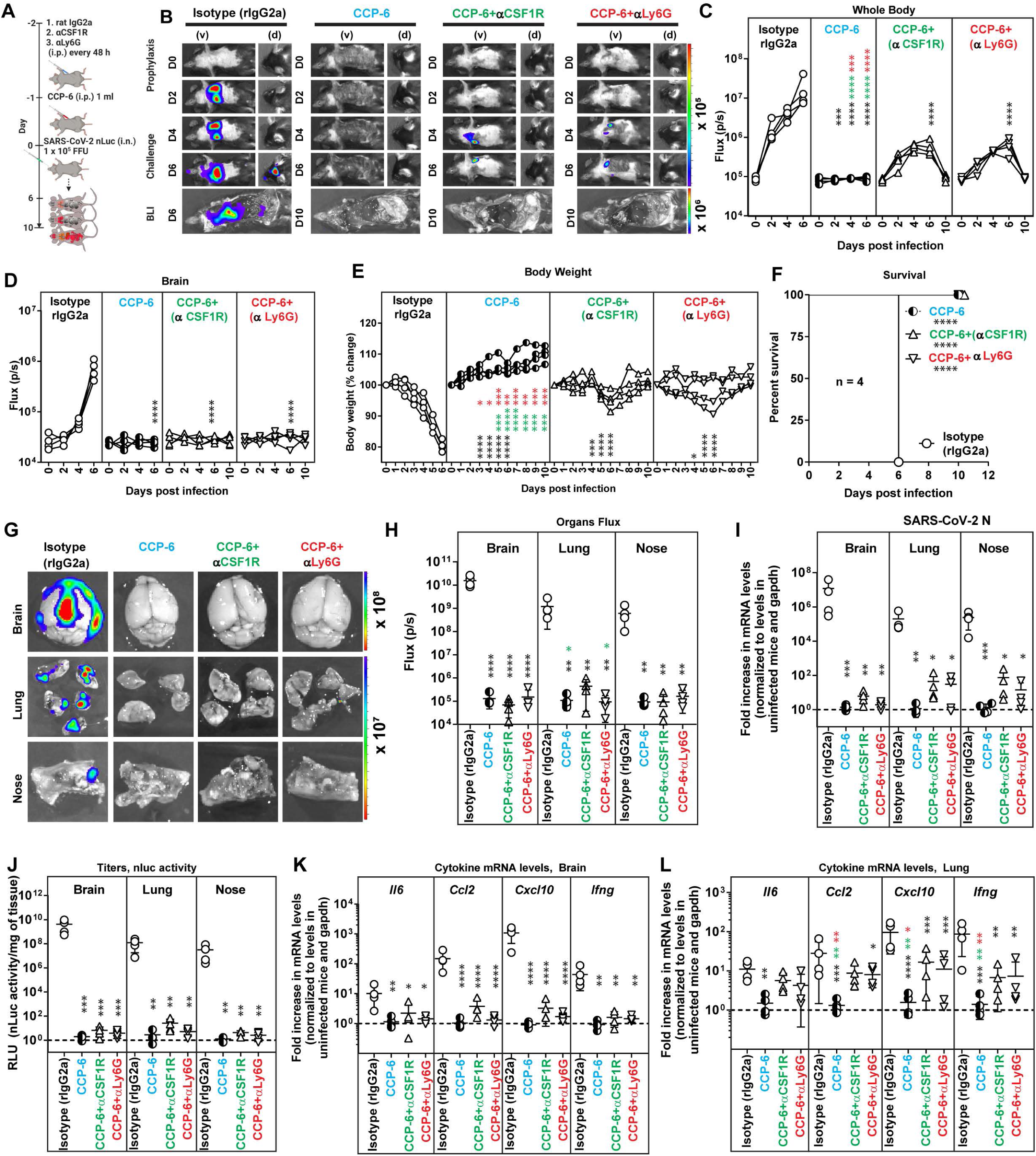
Macrophage And Neutrophil Depletion do not Compromise Protection Against SARS-CoV-2 Infection during CCP Prophylaxis in K18-hACE2 Mice. (A) Experimental design to test the contribution of macrophages (CD45^+^Ly6G^-^Ly6C^-^ CD11b^+^CD68^+^) and neutrophils CD45^+^CD11b^+^Ly6G^+^) in K18-hACE2 mice challenged with SARS-CoV-2-nLuc (1 x 10^5^ FFU, i.n.) and treated prophylactically (i.p.; -1 dpi, 1 ml/ 20-25 g body weight) with CCP-6. αCSFR-1 or αLy6G mAbs (i.p., 20 mg/kg body weight) were used to deplete macrophages and neutrophils respectively every 48h starting two days before infection. Human and rat isotype mAb-treated cohorts served as controls (Iso). Animals were followed by non-invasive BLI every 2 days as indicated. (B) Representative BLI images of SARS-CoV-2-nLuc-infected mice in ventral (v) and dorsal (d) positions. Scale bars denote radiance (photons/sec/cm^2^/steradian). (C-D) Temporal quantification of nLuc signal as flux (photons/sec) computed non-invasively. (E) Temporal changes in mouse body weight with initial body weight set to 100% for an experiment shown in A. (F) Kaplan-Meier survival curves of mice (n = 4 per group) statistically compared by log-rank (Mantel-Cox) test for experiment as in A. (G, H) *Ex vivo* imaging of indicated organs and quantification of nLuc signal as flux(photons/sec) after necropsy. (I) Fold change in SARS-CoV-2 nucleocapsid (N gene) expression in brain, lung and nose tissues. The data were normalized to *Gapdh* mRNA expression in the same sample and that in non-infected mice after necropsy. (J) Viral loads (nLuc activity/mg) in indicated tissue measured on Vero E6 cells as targets. Undetectable virus amounts were set to 1. (K, L) Fold change in cytokine mRNA expression in brain and lung tissues. The data were normalized to *Gapdh* mRNA expression in the same sample and that in non-infected mice after necropsy. Viral loads (I, J) and inflammatory cytokine profile (K, L) were determined after necropsy for mice that succumb to infection at 6 dpi and for surviving mice at 10 dpi. Each curve in C-E and each data point in H-L represents an individual mouse. Data in panels C-L are from two independent experiments and n=2 mouse per group. Grouped data in (C-E), (H-L) were analyzed by 2-way ANOVA followed by Tukey’s multiple comparison tests. Statistical significance for group comparisons to isotype control are shown in black, with CCP-6 are shown in cyan, with CCP-6+αCSF1R are shown in green and with CCP-6 αLy6G are shown in red. ∗, p < 0.05; ∗∗, p < 0.01; ∗∗∗, p < 0.001; ∗∗∗∗, p < 0.0001; Mean values ± SD are depicted. See also Figure S2

### Neutrophils and Macrophages Contribute Significantly to CCP Potency During Therapy

Next, we depleted neutrophils (anti-Ly6G) or macrophages (anti-CSF1R) to analyze the role of innate effector cells toward the therapeutic effects of CCP-6 *in vivo* **(Figure 4A)**. Longitudinal BLI analyses and nLuc signal quantification revealed that depletion of either macrophages or neutrophils significantly compromised CCP-6-mediated virologic control **(Figure 4B-D, Figure S3)**. Innate immune cell-depleted cohorts lost ∼20-30 % of their body weight and succumbed to infection **(Figure 4E, F)** with 50% and 100% of mice in macrophage and neutrophil-depleted cohorts respectively underwent a one-day delay in death upon CCP-6 treatment. Virus neuroinvasion occurred in 50 and 75% of mice (respectively) but after a 2-day delay compared to isotype antibody-treated mice (4 vs 6 dpi) reflecting the ability of CCP-6 to prevent virus neuroinvasion upto an extent even in the absence innate cells. These data suggested a more significant contribution of macrophages compared to neutrophils in CCP-6-mediated Fc effector functions during therapy. Innate immune cell depletion compromised CCP-6-mediated virologic control resulting in higher viral loads in the nose and lungs like control cohorts (hIgG1 and rat IgG2A-treated) at experimental endpoints **(Figure 4G)**. The ability of CCP-6 to prevent exacerbated expression of inflammatory cytokine mRNAs (CCL2, CXCL10, IFNG) in the lungs was also significantly compromised when neutrophils or macrophages were depleted **(Figure 4H, I)**. However, inflammatory cytokines (CCL2, CXCL10) in the brain remained under control reflecting the delay in neuroinvasion compared to isotype antibody-treated cohorts **(Figure 4H)**. These data show that Fc-effector functions mediated by innate immune effector cells significantly contributed to CCP-6-mediated protection during therapy and were also required to dampen inflammation especially in the lungs where SARS-CoV-2 established infection.

**Figure 4.**
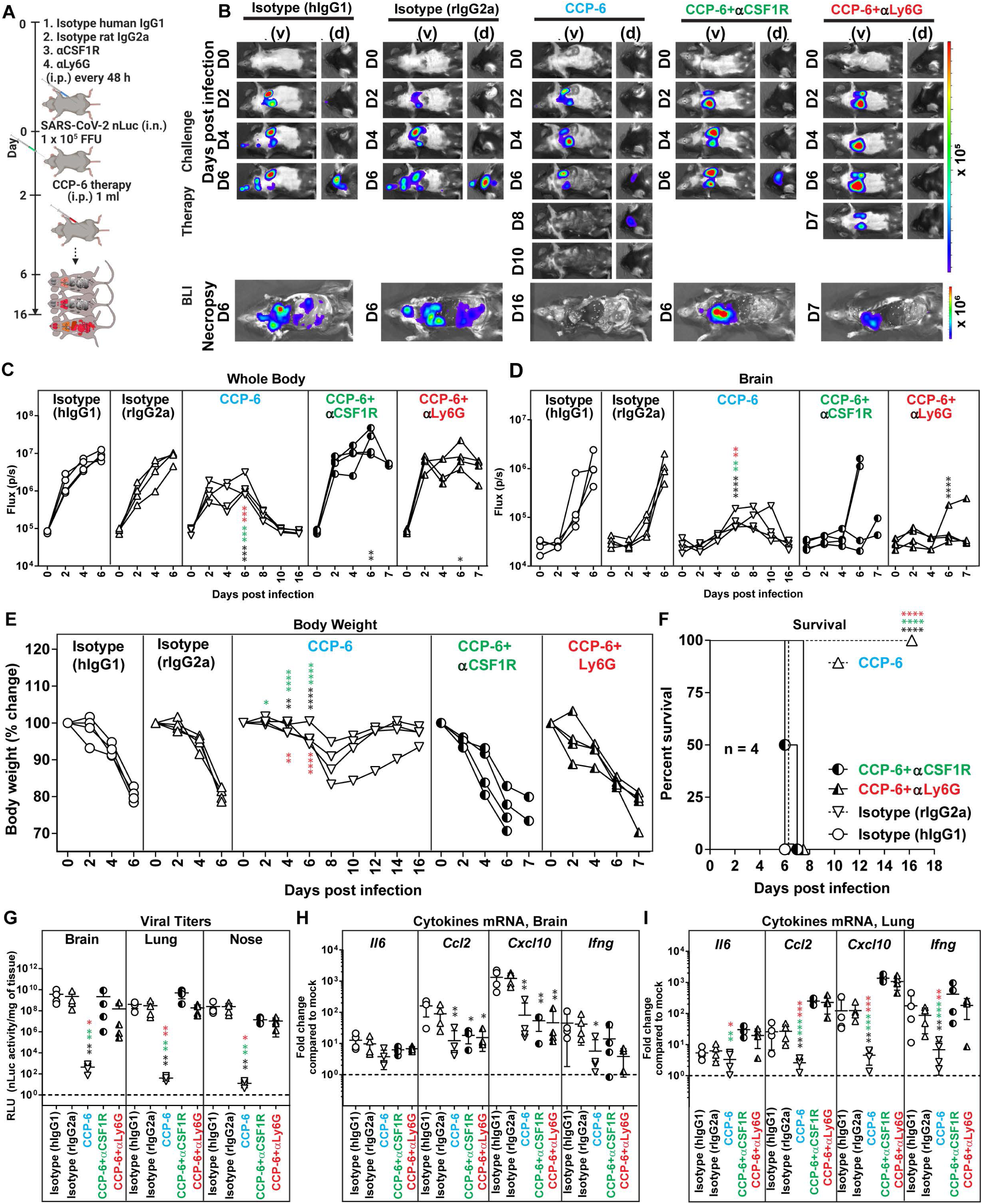
Macrophages and Neutrophils Are Required to Eliminate Established SARS-CoV-2 Infection During CCP Therapy in K18-hACE2 Mice. (A) Experimental design to test the contribution of macrophages (CD45^+^CD11b^+^CD68^+^) and neutrophils (CD45^+^CD11b^+^Ly6G^+^) in K18-hACE2 mice therapeutically treated at 2 dpi with CCP-6 (i.p., 1 ml/ 20-25 g body weight) after challenge with SARS-CoV-2-nLuc (i.n., 1 x 10^5^ FFU). αCSF1R or αLy6G mAbs (i.p., 20 mg/kg body weight) were used to deplete macrophages and neutrophils respectively every 48h starting at 0 dpi. Human and/or rat isotype mAb treated cohorts served as controls (Isotype). The mice were followed by non-invasive BLI every 2 days from the start of infection. (B) Representative BLI images of SARS-CoV-2-nLuc-infected mice in ventral (v) and dorsal (d) positions. Scale bars denote radiance (photons/sec/cm^2^/steradian). (C-D) Temporal quantification of nLuc signal as flux (photons/sec) computed non-invasively. (E) Temporal changes in mouse body weight with initial body weight set to 100% for an experiment shown in A. (F) Kaplan-Meier survival curves of mice (n = 4 per group) statistically compared by log-rank (Mantel-Cox) test for experiment as in A. (G) Viral loads (nLuc activity/mg) from indicated tissue using Vero E6 cells as targets. Undetectable virus amounts were set to 1. (H, I) Fold change in cytokine mRNA expression in brain and lung tissues. The data were normalized to *Gapdh* mRNA expression in the same sample and that in non-infected mice after necropsy. Viral loads (G) and inflammatory cytokine profile (H, I) were determined at 6 dpi for mice that succumbed to infection and for surviving mice at 10 dpi after necropsy. Each curve in (C-E) and each data point in (G-I) represents an individual mouse. Data in panels C-I are from two independent experiments and n=2-3 mice per group. Grouped data in (C-E), (G-I) were analyzed by 2-way ANOVA followed by Tukey’s multiple comparison tests. Statistical significance for group comparisons to isotype control are shown in black, with CCP-6-treated cohorts are shown as cyan, CCP-6-treated neutrophil-depleted cohorts are shown in red, and with CCP-treated macrophage-depleted cohorts are shown in green. ∗, p < 0.05; ∗∗, p < 0.01; ∗∗∗, p < 0.001; ∗∗∗∗, p < 0.0001; Mean values ± SD are depicted. See also Figure S3.

### Polyconal IgGs Contribute to Protection During CCP Prophylaxis

IgM and IgA are mucosal antibodies that function as the first line of defense against mucosal pathogens (Russell et al., 2020). Although not as potent as IgG, multivalent antibodies like IgM (pentamer: decavalent) and IgA (dimer: tetravalent) can exhibit enhanced neutralization due to their avidity (Gasser et al., 2021; Ku et al., 2021; Wang et al., 2021). We depleted IgG or Ig(M+A) from CCP-6 to evaluate the contribution of specific antibody classes towards protection. We confirmed successful depletion of antibody class-subsets by class-specific Ig ELISA [<99% of IgG or 90-95% of Ig(M+A)] **(Figure S4A)** and flow-cytometric evaluation of Spike-specific isotype content using Spike-expressing HEK293 cells **(Figure S4B)**. ADCC analyses of the undepleted and depleted CCP-6 revealed that *in vitro* Fc- activities predominantly tracked with Ig(M+A)- depleted fraction **(Figure S4C)**. While both fractions displayed SARS-CoV-2 neutralizing activity **(Figure S4D)**, the Ig(M+A) depleted fraction (containing IgG) demonstrated ∼2.3-fold higher neutralizing activity than the IgG-depleted-fraction.

We next investigated the anti-SARS-CoV-2 *in vivo* efficacy of class-depleted plasma fractions during prophylaxis **(Figure 5A)**. Unfractionated CCP-6 was diluted before use to account for the loss in IgG (Equalized IgG) in the Ig(M+A)-depleted fraction incurred during the depletion procedure. Longitudinal BLI revealed that IgG-depletion led to a near-complete loss in CCP-6 mediated protection with uncontrolled virus replication, neuroinvasion, 15-20% body weight loss and 100% mortality **(Figure 5B-F, Figure S5A-B)**. In contrast, Ig(M+A)-depleted fraction displayed virologic control like undepleted CCP-6 (Equalized IgG) with 100% survival efficacy **(Figure 5F)** despite a small reduction (<10%) in body weight compared to undepleted plasma **(Figure 5E, F)**. Significantly higher viral loads and inflammatory cytokine mRNA expression in target organs reflected the loss of virologic control in mice treated with IgG-depleted CCP-6 compared to mice treated with unfractionated and Ig(M+A)-depleted plasma **(Figure 5G-I)**. Thus, polyclonal IgGs predominantly contributed to virologic control and protection with marginal but distinct contribution from polyclonal Ig(M+A) during CCP-6 prophylaxis.

**Figure 5.**
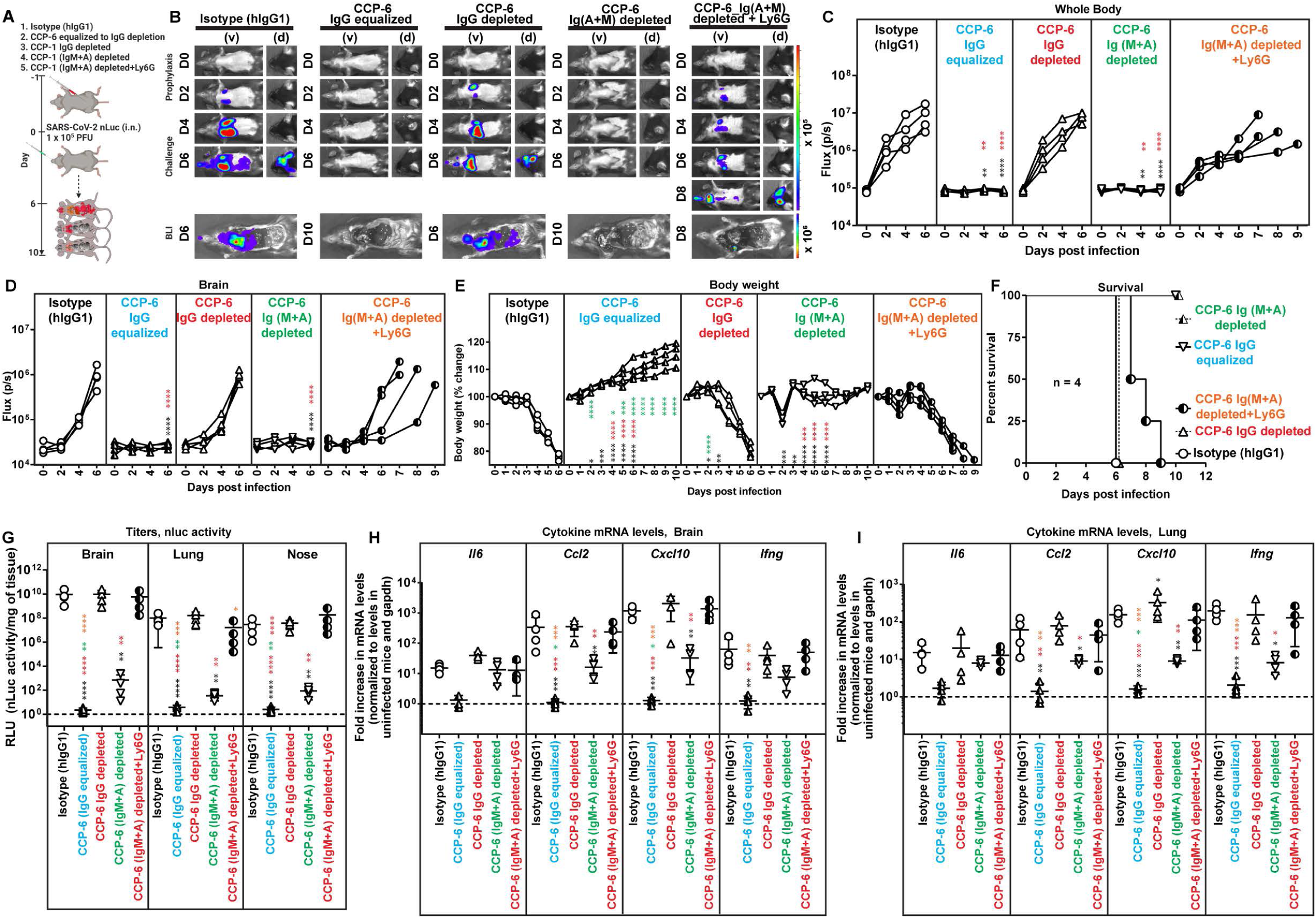
Polyclonal IgGs in CCP Predominantly Contribute to Protection During Prophylaxis in SARS-CoV-2-infected K18-hACE2 Mice. (A) Experimental design to test *in vivo* efficacies of IgG- and Ig(M+A)-depleted CCP-6 (1 ml x 2 i.p. injections, 3 h apart) in SARS-CoV-2-nLuc infected K18-hACE2 mice (i.n., 1 x10^5^ FFU) under prophylaxis (-1 dpi). For CCP-6 treatment, plasma was diluted to equalize IgG content of Ig(M+A)- depleted plasma. αLy6G mAb (i.p., 20 mg/kg body weight) was used to deplete neutrophils respectively every 48h starting two days before infection. Mice treated with hIgG1 served as controls (Iso). The mice were followed by non-invasive BLI every 2 days from the start of infection. (B) Representative BLI images of SARS-CoV-2-nLuc-infected mice in ventral (v) and dorsal (d) positions. Scale bars denote radiance (photons/sec/cm^2^/steradian). (C-D) Temporal quantification of nLuc signal as flux (photons/sec) computed non-invasively. (E) Temporal changes in mouse body weight with initial body weight set to 100% for an experiment shown in A. (F) Kaplan-Meier survival curves of mice (n = 4 per group) statistically compared by log-rank (Mantel-Cox) test for experiment as in A. (G) Viral loads (nLuc activity/mg) from indicated tissues using Vero E6 cells as targets. Undetectable virus amounts were set to 1. (H, I) Fold change in cytokine mRNA expression in brain and lung tissues. The data were normalized to *Gapdh* mRNA expression in the same sample and that in non-infected mice after necropsy. Viral loads (G) and inflammatory cytokine profile (H, I) were determined at the time of death at 6 dpi or 10 dpi for surviving mice after necropsy. Each curve in C-E and each data point in G-I represents an individual mouse. Data in panels C-I are from are from two independent experiments and n=2 mice per group Grouped data in (C-E), (G-I) were analyzed by 2-way ANOVA followed by Tukey’s multiple comparison tests. Statistical significance for group comparisons to isotype control are shown in black, with IgG equalized CCP-6 are shown in cyan, with IgG-depleted CCP-6 are shown in red, with Ig(M+A)-depleted CCP-6 are shown in green and with Ig(M+A)-depleted CCP under neutrophil depletion are shown in orange. ∗, p < 0.05; ∗∗, p < 0.01; ∗∗∗, p < 0.001; ∗∗∗∗, p < 0.0001; Mean values ± SD are depicted. See also Figure S4.

To decipher if direct neutralization and/or Fc-mediated innate cell-recruitment contributed to protection during prophylaxis with Ig(M+A)-depleted plasma (containing IgG), we immuno-depleted neutrophils (anti-Ly6G). Interestingly, compared to the undepleted plasma where innate cells contributed marginally during prophylaxis, neutrophil depletion had a significant impact on protection conferred by Ig(M+A)-depleted fraction **(Figure 5B-F)**. BLI analyses revealed loss of virologic control with visible infection at 2-4 dpi and dissemination of virus into the brain at 8 dpi **(Figure 5B, D S5A, B)** with all the mice in the neutrophil-depleted cohort losing weight and succumbing to SARS-CoV-2 challenge, albeit with a delay of 1-3 days **(Figure 5E, F)**. These data correlated with increased viral loads in tissues and enhanced inflammatory cytokine mRNA expression in neutrophil-depleted cohorts prophylactically treated with Ig(M+A)-depleted CCP **(Figure 5G-I, S5C)**. These data suggest a functional interplay between Ig(M+A) and IgG to promote virus neutralization. When Ig(M+A) was depleted, the reliance on Fc-effector mechanisms over direct neutralization by the IgG-harboring fraction was significantly increased for effective virological control and was largely mediated by neutrophils. Thus, when neutralization potency in Ig(M+A) depleted CCP-6 was insufficient to prevent virus infection, IgG-driven Fc-effector recruitment of innate immune cells acted as a second line of defense to promote infected-cell clearance and control virus replication during prophylaxis.

### Polyclonal IgG and Ig(M+A) Fc-effector Activities Are Required for *In Vivo* CCP Efficacy During Therapy

Longitudinal BLI analyses revealed that the *in vivo* efficacy of both IgG and Ig(M+A)-depleted plasma against SARS-CoV-2 was severely compromised compared to undepleted CCP-6 during therapy **(Figure 6A-D)**. SARS-CoV-2-nLuc replicated and disseminated to the brain in 6 out of 7 mice in both cohorts that received Ig class-depleted plasma **(Figure 6B-D, G, S6)**. Although 14% of the mice (1 out of 7) in both cohorts survived, body weight and survival analyses showed that mice that received Ig(M+A)-depleted plasma exhibited decelerated body weight loss and delayed mortality (6 dpi vs 8 dpi) compared to those with IgG-depleted plasma **(Figure 6E, F)**. These data suggested that the contribution of IgG was marginally higher than Ig(M+A) to CCP-6 mediated protection. The capacity of CCP-6 to inhibit tissue virus replication and inflammation was also significantly compromised compared to mice treated with undepleted plasma **(Figure 6G-I)**. Interestingly, cytokine mRNA expression (*CCL2 and CXCL10)* in the lungs of mice that received depleted plasma fractions were significantly higher than the unfractionated CCP-6-treated and the isotype IgG1-treated cohorts (Figure 6I). These data reveal the contribution of both immunoglobulin fractions in dampening exacerbated inflammatory immune responses. Thus, as with prophylaxis, Ig class-depletion analyses suggest a functional interplay between IgG and Ig(M+A) for optimal *in vivo* efficacy of CCP.

**Figure 6.**
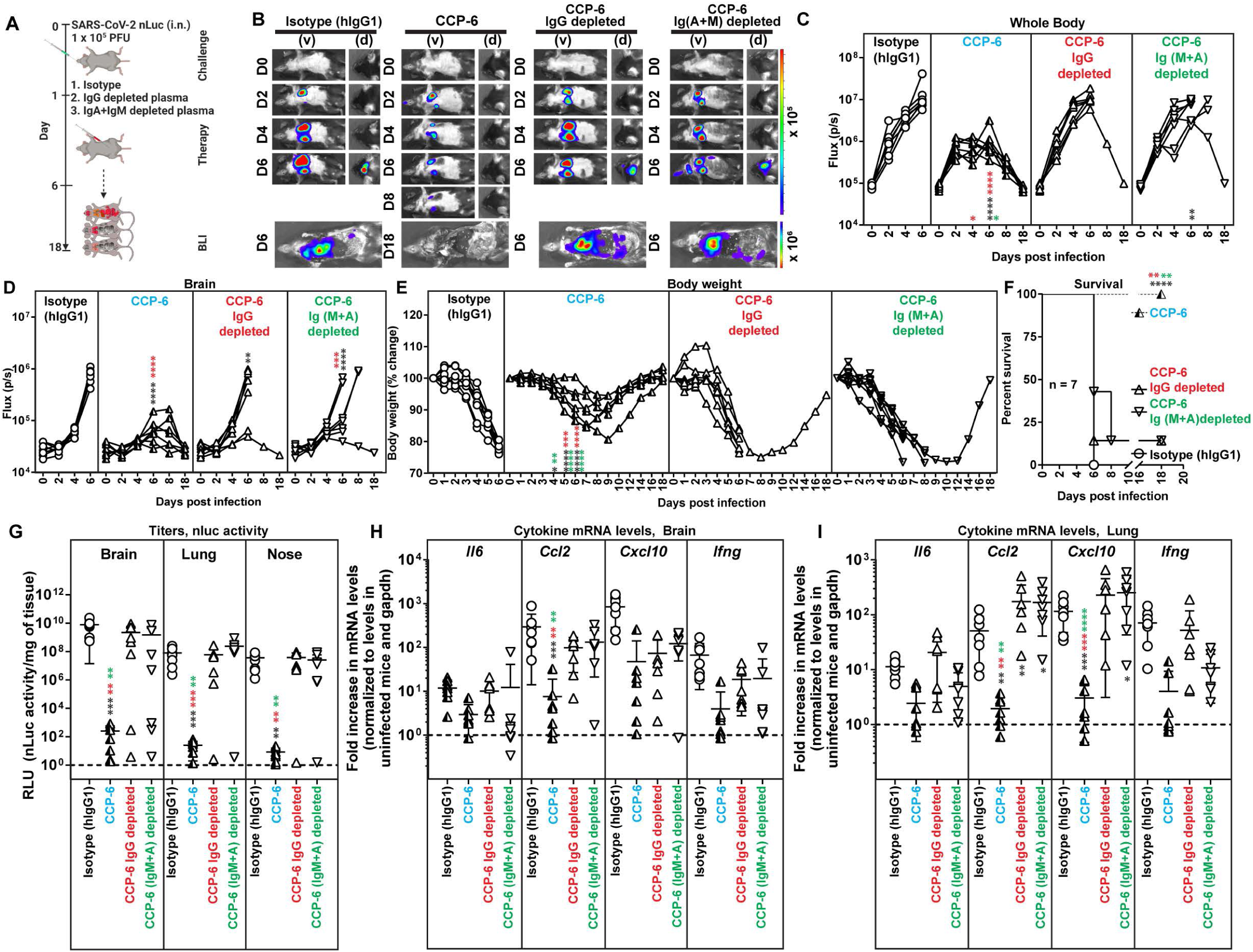
Antibody Classes Collaborate to Achieve Maximal *In Vivo* Protection during CCP Therapy in SARS-CoV-2-infected K18-hACE2 Mice. (A) Experimental design to test *in vivo* efficacies of IgG- and Ig(M+A)-depleted CCP-6 (1 ml x 2 i.p. injections, 1 h apart) in SARS-CoV-2-nLuc infected mice K18-hACE2 mice (i.n., 1 x10^5^ FFU) under therapy (+2 dpi). For CCP-6 treatment, plasma was diluted to equalize IgG content of Ig(M+A)-depleted plasma. Mice treated with hIgG1 served as controls (Iso). The mice were followed by non-invasive BLI every 2 days from the start of infection. (B) Representative BLI images of SARS-CoV-2-nLuc-infected mice in ventral (v) and dorsal (d) positions. Scale bars denote radiance (photons/sec/cm^2^/steradian). (C-D) Temporal quantification of nLuc signal as flux (photons/sec) computed non-invasively. (E) Temporal changes in mouse body weight with initial body weight set to 100% for an experiment shown in A. (F) Kaplan-Meier survival curves of mice (n = 7 per group) statistically compared by log-rank (Mantel-Cox) test for experiment as in A. (G) Viral loads (FFUs/mg) from indicated tissue using Vero E6 cells as targets. Undetectable virus amounts were set to 1. (H, I) Fold change in cytokine mRNA expression in brain and lung tissues. The data were normalized to *Gapdh* mRNA expression in the same sample and that in non-infected mice after necropsy. Viral loads (G) and inflammatory cytokine profile (H, I) were determined at the time of death for mice that succumbed to infection (F) and at 18 dpi for surviving mice Each curve in C-E and each data point in G-I represents an individual mouse. Data in panels C-I are from from two to three independent experiments n=2-3 mouse per group. Grouped data in (C-E), (G-I) were analyzed by 2-way ANOVA followed by Tukey’s multiple comparison tests. Statistical significance for group comparisons to isotype control are shown in black, with IgG-equated CCP-6 are shown in cyan, with IgG-depleted CCP-6 are shown in red and with Ig(M+A)-depleted CCP-6 are shown in green. ∗, p < 0.05; ∗∗, p < 0.01; ∗∗∗, p < 0.001; ∗∗∗∗, p < 0.0001; Mean values ± SD are depicted. See also Figure S6.

### VOC Cross-neutralizing Activity in CCPs is Crucial for protection Against VOCs

Recent *in vitro* studies suggest that broad Fc-effector functions elicited by prior infection or vaccination may offer continued protection against emerging variants of concerns (VOCs) despite loss in neutralization (Kaplonek et al., 2022a; Kaplonek et al., 2022b; Richardson et al., 2022). However, if cross-reactive Fc effector functions can provide *in vivo* protective efficacy when neutralization is diminished remains unexplored. We sought to extend these observations to *in vivo* studies using Wuhan-elicited CCPs against heterologous SARS-CoV-2 VOCs B.1.617.2 (Delta) and B.1.352 (Beta). A comparison of neutralizing IC_50_ values using live virus revealed that except for CCP-1 and CCP-5, all other CCPs had comparable neutralizing activities against both WA1 and Delta VOC. In contrast, the ability to neutralize Beta VOC compared to WA1 was significantly diminished for all the CCPs tested **(Figure 7A)**.

**Figure 7.**
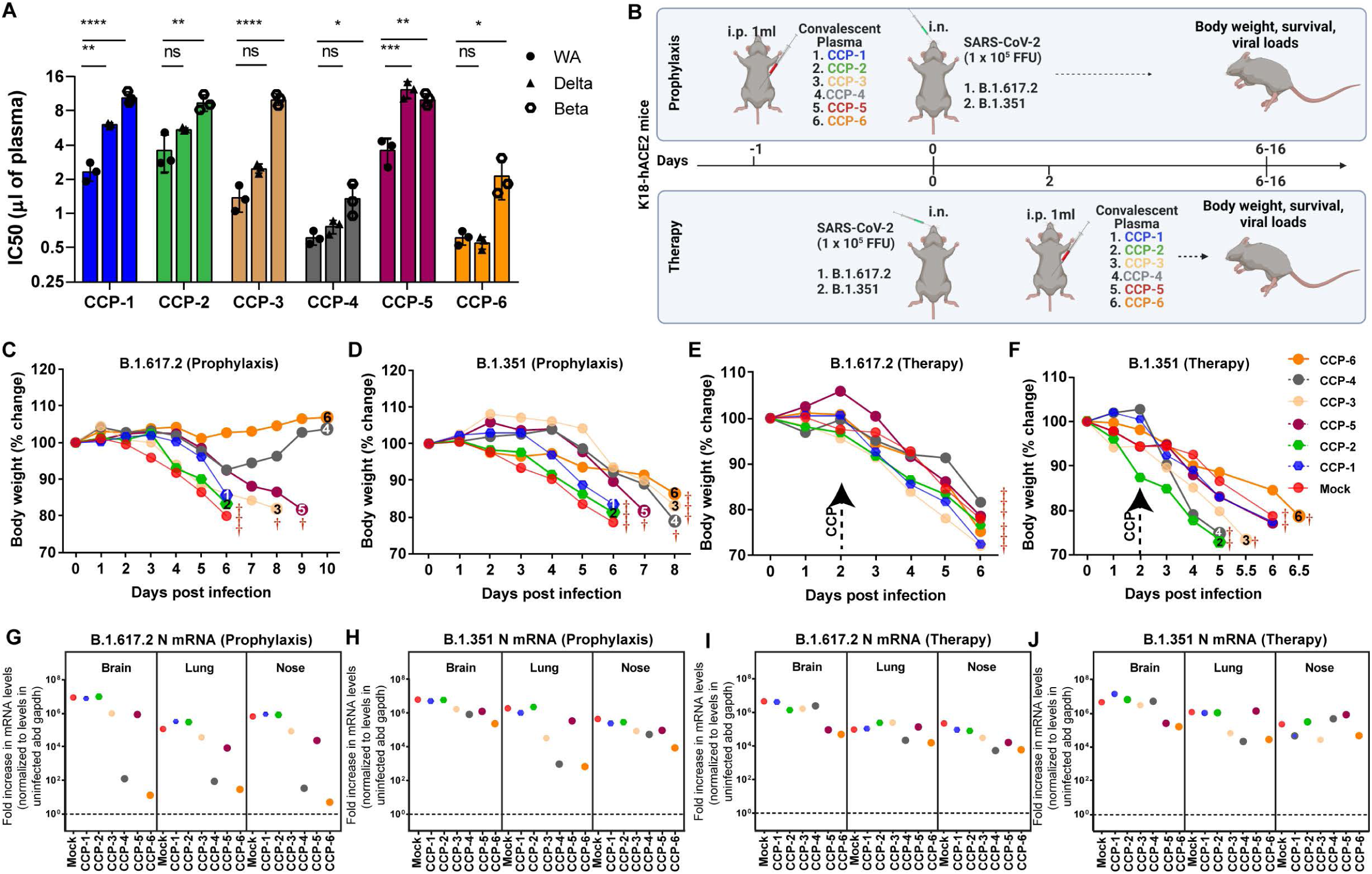
Fc-mediated cross-protective efficacy profiles of WA1-elicited CCPs against Delta and Beta VOCs in K18-hACE2 mice. (A) A graph depicting WA1, Delta and Beta-neutralizing activity of indicated CCPs expressed as inhibitory concentration of plasma (IC_50_) that reduces FFUs by 50% using Vero E6 cells as targets. One-way Anova with Dunnett’s multiple comparisons test was used to determine is if Delta and Beta VOC neutralizing titers in CCPs differed significantly to WA1-neutralizing titers. p < 0.05; ∗∗, p < 0.01; ∗∗∗, p < 0.001; ∗∗∗∗, p < 0.0001; Mean values ± SD are depicted. (B) A scheme showing experimental design for screening *in vivo* efficacy of indicated CCPs delivered 1ml per 20-25 g body weight of mouse intraperitoneally (i.p.) under prophylaxis (-1dpi) and therapeutically (+2 dpi) in K18-hACE2 mice intranasally (i.n.) challenged with 1 x 10^5^ FFU of B.1.617.2 (Delta VOC) or B.1.351 (Beta VOC). PBS-treated mice were used as control (Mock). (C-F) Temporal changes in mouse body weight with initial body weight set to 100% during CCP prophylaxis (-1dpi) and therapy (+2 dpi) for experiment as in C in mice challenged with Delta and Beta VOC. (G-J) Fold change in SARS-CoV-2 nucleocapsid (N gene) expression in brain, lung and nose tissues during CCP prophylaxis and therapy for experiment shown in C. The data were normalized to *Gapdh* mRNA expression in the same sample and that in non-infected mice after necropsy.

We next examined the *in vivo* efficacy of CCPs in K18-hACE2 mice challenged with Delta and Beta VOCs under prophylaxis (-1dpi) and therapy (+2 dpi) **(Figure 7C)**. Prophylaxis using CCP-1 and CCP-2 with low Fc-effector activities failed to protect against both VOCs and the mice exhibited body weight loss and death at 6 dpi like mock-treated control (**Figure 7C, D)**. In contrast, prophylaxis with CCP-4 and CCP-6, that retained considerable Delta VOC-neutralizing activity (IC50 <1/X of plasma) protected mice against Delta and prevented mortality. Prophylaxis with CCP-3 and CCP-5 delayed mortality by 2 and 3 days respectively. Notably, CCP-5, with the weakest neutralizing activity against Delta, delayed mortality which suggested a contribution of Fc-functions towards VOC immunity. These data correlated with lower N mRNA expression in target organs for CCP-4 and 6 followed by CCP-5 and 3-treated mice compared to mock-treated mouse **(Figure 7G, H)**. None of the CCPs analyzed protected against Beta VOC-induced mortality under prophylaxis **(Figure 7D)**. However, treatment with CCP-3, 4, 5 and 6 delayed Beta VOC- induced death by 1-2 days and lowered lung viral loads compared to mock and CCP-1 and 2 treated animals **(Figure 7D, H)**. While existing Beta-neutralizing activity likely explained the delayed mortality and reduction in viral loads in CCP-4 and 6 treated mice, Fc-effector functions may have contributed to the marginal protection seen in CCP-3 and 5-treated mice. Notably, none of the CCPs prevented Delta or Beta VOCs-induced mortality under therapy (+2 dpi) **(Figure 7E, F)**. We did observe half a day delay in death during Beta VOC infection in mouse under CCP-6 therapy. In addition, therapy with CCP-4 and 6 lowered lung viral loads for both Delta and Beta VOC **(Figure 7I, J)**. Thus, while cross-VOC Fc-effector functions contributed to immunity, they were insufficient to prevent VOC-induced mortality in mice and VOC-neutralizing activity in CCPs remained vital for protection against VOCs.

## Discussion

The constituents of convalescent plasma are complex, and it is difficult to predict their *in vivo* efficacies based solely on neutralizing titers or Spike-specific immunoglobulin content. To navigate the intricacies of CCPs, additional measures of selection that track with *in vivo* protection are required and important to guide best practices in future infectious disease outbreaks. Furthermore *in vivo* models that allow testing CPs with protective profiles can help identify suitable properties that can be incorporated in high throughput screening assays *in vitro*. Here we combined the highly susceptible K18-hACE2 mouse model of SARS-CoV-2 with bioluminescence imaging to track virus replication for studying efficacies and characteristics of CCPs that contribute to *in vivo* protection. We show that CCPs with robust Fc effector activities, but low neutralization can protect against homologous SARS-CoV-2 infection under both therapy and prophylaxis. Antibody isotypes, IgG, IgM and IgA collaborated for optimal virologic control and polyclonal Fc-mediated innate immune cell recruitment was required for effective clearing of established SARS-CoV-2 infection. Additionally, despite reduced neutralization, CCPs provided some levels of cross-immunity against Delta and Beta VOCs likely via Fc-mediated effector activities by delaying mortality in mice. Thus, our study underscores the utility of *in vivo* models combined with live tracking of virus replication to identify suitable properties of CCPs with net protective profiles and revealed the contribution of polyclonal Fc-effector functions and antibody classes in combating SARS-CoV-2 and VOCs.

We chose CCPs with low neutralizing activities (ID_50_ ≤1:250) together with immune cell depletion to evaluate the *in vivo* efficacy of associated Fc-effector activities. Polyclonal neutralizing activity present in CCP-6 was sufficient to protect mice from SARS-CoV-2-induced mortality during prophylaxis. Despite neutralization playing a major role during CCP-6 prophylaxis, visible infection in the lungs together with transient weight loss upon depletion of innate immune cells revealed a minor yet distinct contribution of Fc-effector functions towards protection. The contribution of Fc-effector mechanism to protection increased significantly when Ig(M+A) was depleted from CCP-6. The combined effect of IgG-driven neutralizing activity and polyclonal Fc-effector functions was however able to compensate for lost Ig(M+A) associated activities to protect mice from SARS-CoV-2. These data also revealed that Ig(M+A) collaborated with IgG to potentiate overall neutralizing activity. A sufficiently potent neutralizing activity alone can counter incoming free virus during SARS-CoV-2 challenge under prophylaxis. However, during therapy, CCPs need to counter both free virus and infected producer cells for curbing virus replication and spread. Thus, in addition to neutralizing activity, for a CCP to be effective therapeutically, polyclonal Fc-mediated innate cell recruitment is likely paramount for clearing pools of infected cells through mechanisms such as ADCC and ADCP. Indeed, we found a critical requirement for neutrophils and macrophages during CCP therapy. Furthermore, both IgG and Ig(M+A) fractions were required during therapy as depleting either, compromised protection. Like prophylaxis, a collaboration between all antibody isotypes was required for effective virologic control during therapy. Exacerbated inflammatory response is one of the hallmarks of SARS-CoV-2-induced disease. In addition to virologic control, we also found that recruitment of innate immune cells through polyclonal Fc-FcR interaction can dampen SARS-CoV-2-induced inflammatory response. Thus CCP-associated polyclonal Fc-effector functions have the potential to mitigate SARS-CoV-2-induced disease newly infected COVID-19 patients.

CCP Fc-effector function was rarely measured in COVID-19 clinical studies. Given that the CONCOR-1 trial reported only a loose correlation between neutralizing and Fc-effector functions, one can assume that even studies using stringent CCP selection criteria likely used plasma with variable degrees Fc-effector function (Begin et al., 2021). Developing the capacity to adapt and disseminate Fc-effector function testing rapidly may be key to its wider use in future pandemics and a more optimal use of CCP, directing those with high neutralizing but low Fc-effector function toward prophylaxis trials while reserving those with both high neutralizing and high Fc-effector functions for the acutely ill.

The complexities associated with CCP therapy can be resolved using monoclonal antibody cocktails for therapeutic applications (Liu and Shameem, 2022). However, antibody cocktails remain expensive to implement especially in developing nations (Ledford, 2020). They are also subject to neutralization escape by evolving VOCs rendering them ineffective as has been the case with Omicron and its sublineages (Cao et al., 2022; Liu et al., 2022; Planas et al., 2022). In contrast, the ensemble of both nAbs and non-nAbs that CCPs possess make them more resilient to VOCs despite drop in neutralizing titers. Fc-effector mechanisms garnered by SARS-CoV-2-binding antibodies is suggested to remain effective against VOCs (Kaplonek et al., 2022a; Kaplonek et al., 2022b; Richardson et al., 2022) . Whether VOC cross-reactive Fc-effector activities offer *in vivo* protection remains unexplored. Our *in vivo* efficacy analyses in mice under prophylaxis revealed that Fc-effector activities elicited by the ancestral SARS-CoV-2 that caused the first wave can be effective in delaying death under VOC challenge but were not enough to provide complete protection. These data mirrored our previous analyses where a Fc-enhanced non-nAb did not offer complete protection but delayed mortality in mice (Beaudoin-Bussieres et al., 2022). However, combining Fc-enhanced non-nAb with Fc-compromised nAb completely protected mice when each antibody failed to protect on their own. Thus, cross-reactive Fc-effector functions on their own is likely not enough for protection against VOCs. Polyclonal neutralizing activity, although diminished, form a critical component of the mix with Fc-effector activities to engender protection against VOCs. Overall, our *in vivo* analyses endorse inclusion of Fc-effector activities in addition to neutralization as additional criteria to select CCPs for therapeutic applications. Several high throughput *in vitro* assays exist that can examine multiple signatures of CCPs (Begin et al., 2021; Gunn et al., 2021). *In vivo* efficacy analyses could complement these assays to navigate complex CCP characteristics for identifying those with net protective profiles. Identifying plasma signatures that track with protective or detrimental effects will be key to the success of CP therapy for future infectious disease outbreaks and pandemics.

## Author contributions

Conceptualization, PDU, AF, RB and PB; Methodology, PDU, IU, MC, AF and RB; Investigation, IU, KS, PDU, GBB, ED, AT, AL; Writing – Original Draft, PDU; Writing – Review & Editing, PDU, AF, WM, PK, RB, PB, MC, IU; Funding Acquisition, WM, AF, PB, and RB, Resources, WM, PK, AF and RB Supervision, PDU, AF and RB

## Acknowledgements

This work was supported by This work was supported by NIH grant R01AI163395 to WM ,le Ministère de l’Économie et de l’Innovation du Québec, Programme de soutien aux organismes de recherche et d’innovation, Foundation du CHUM, CIHR grant nos. 352417 and 177958, a CFI grant, 41027 and a Canada Research Chair on Retroviral Entry no. RCHS0235 950-232424 to AF; Canada’s COVID-19 Immunity Task Force (CITF) & Canada Foundation for Innovation (CFI) #41027 to AF, CIHR fellowships to GBB, le Ministère de l’Économie et de l’Innovation du Québec, Fondation du CHU Sainte-Justine Fonds de recherché du Québec – Santé #281662 to PB.

## STAR Methods

### KEY RESOURCES TABLE

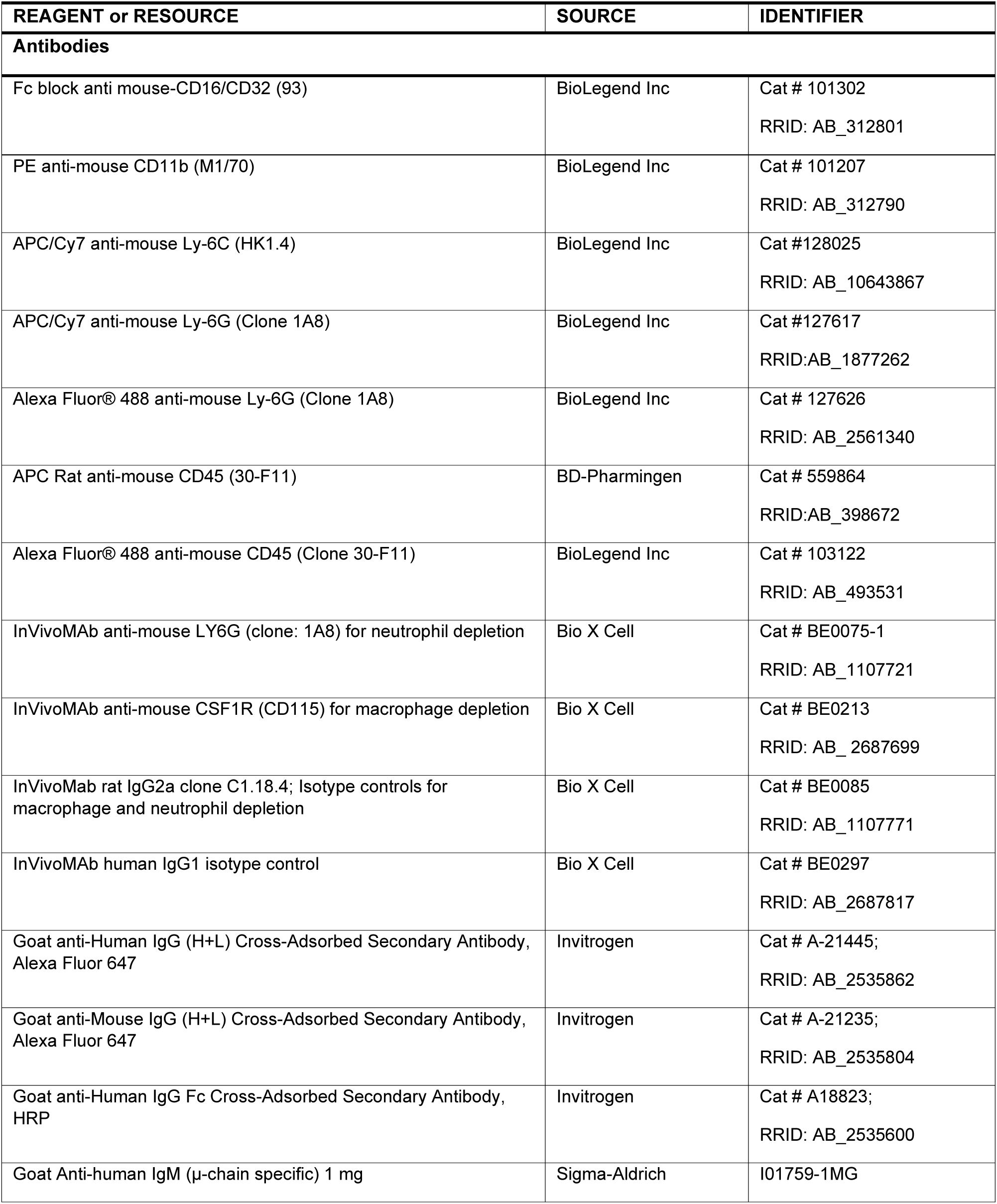
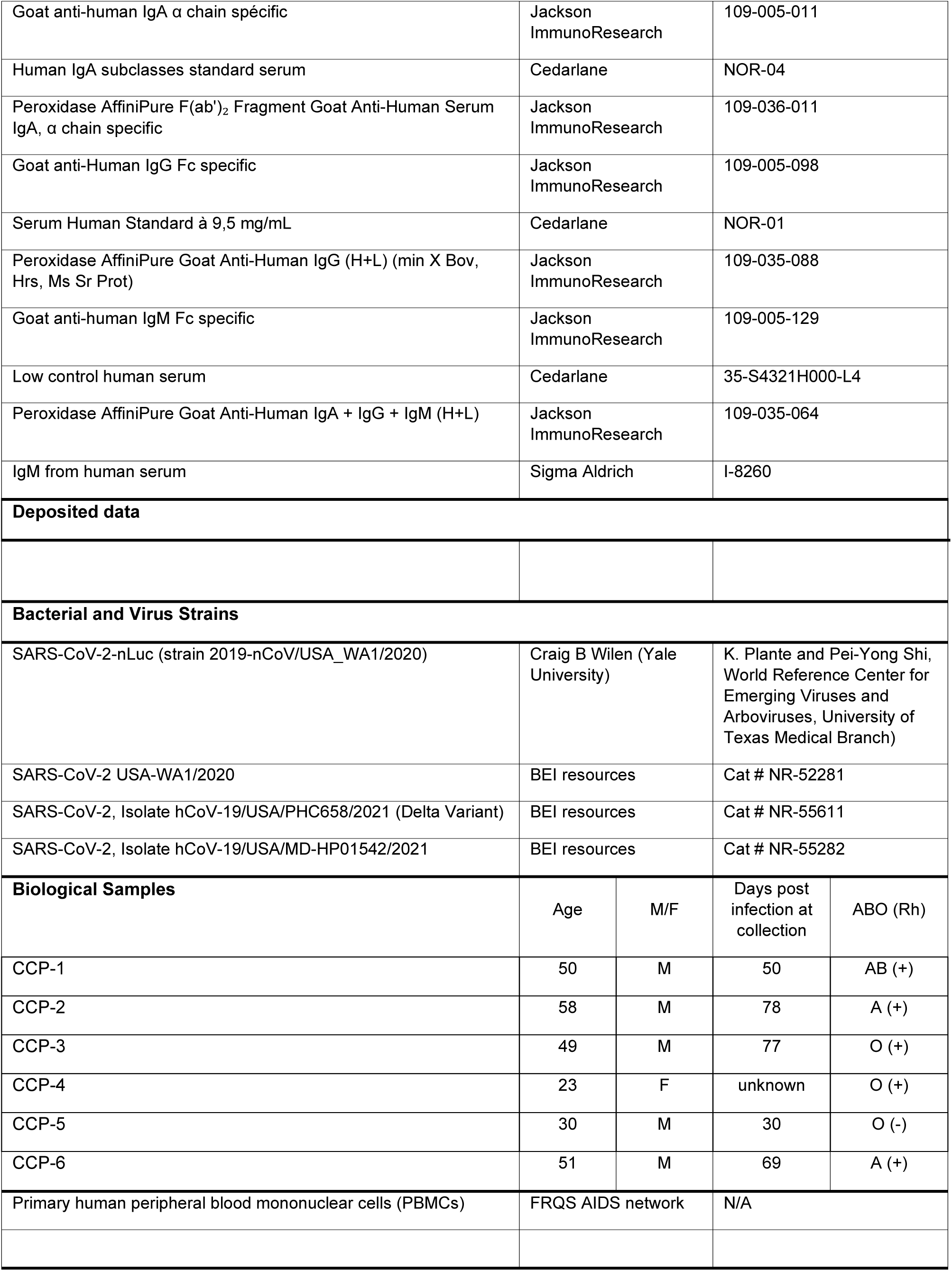
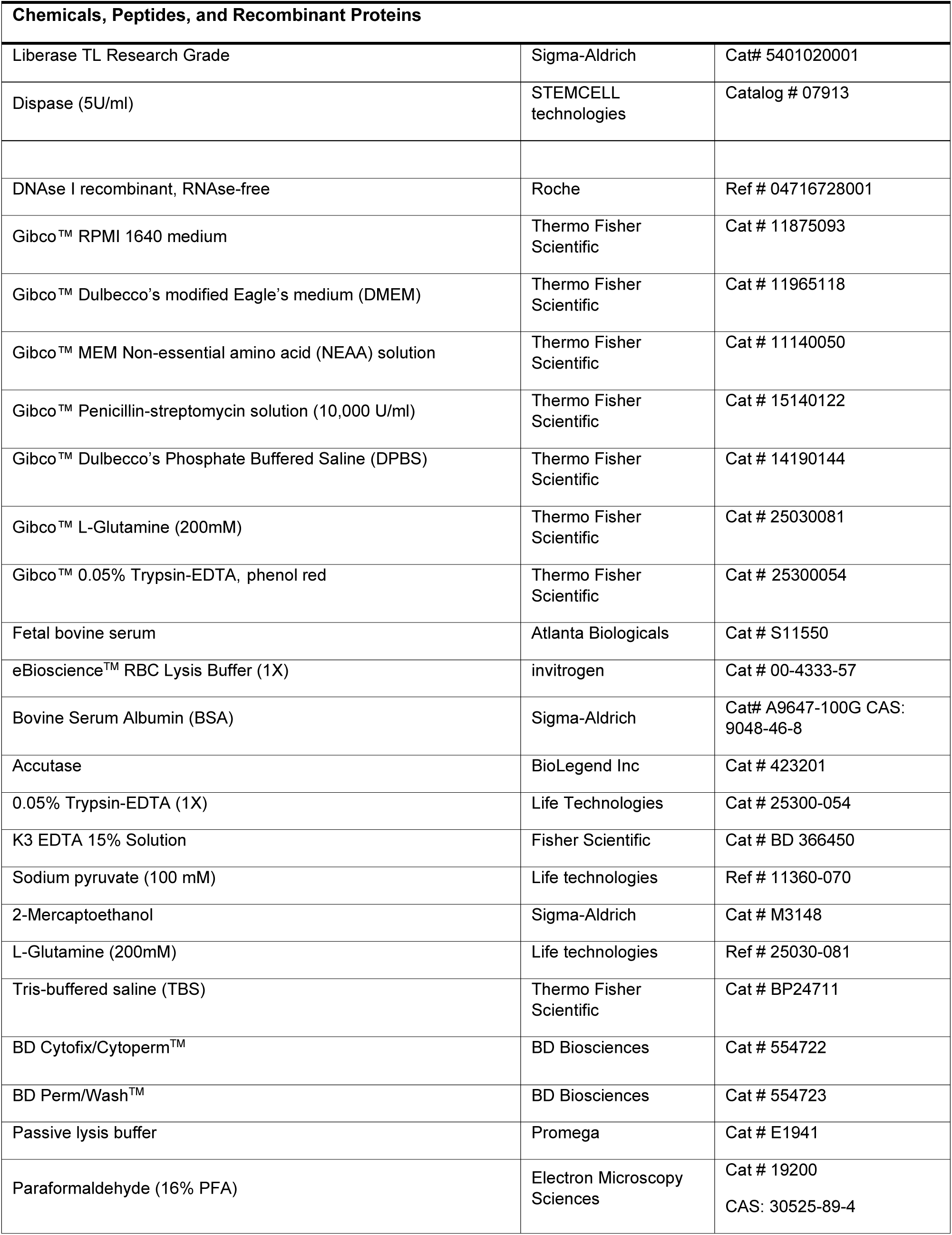
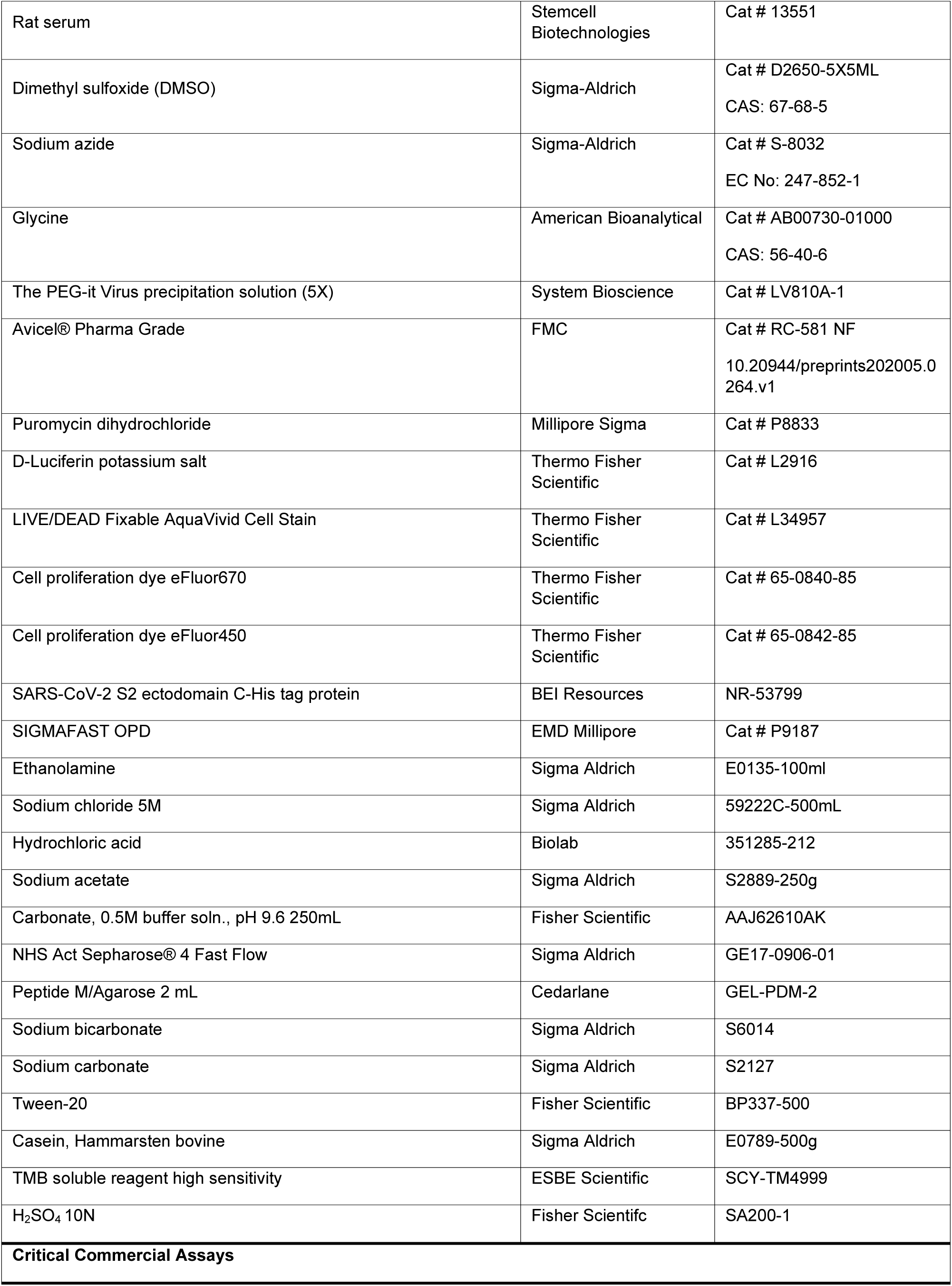
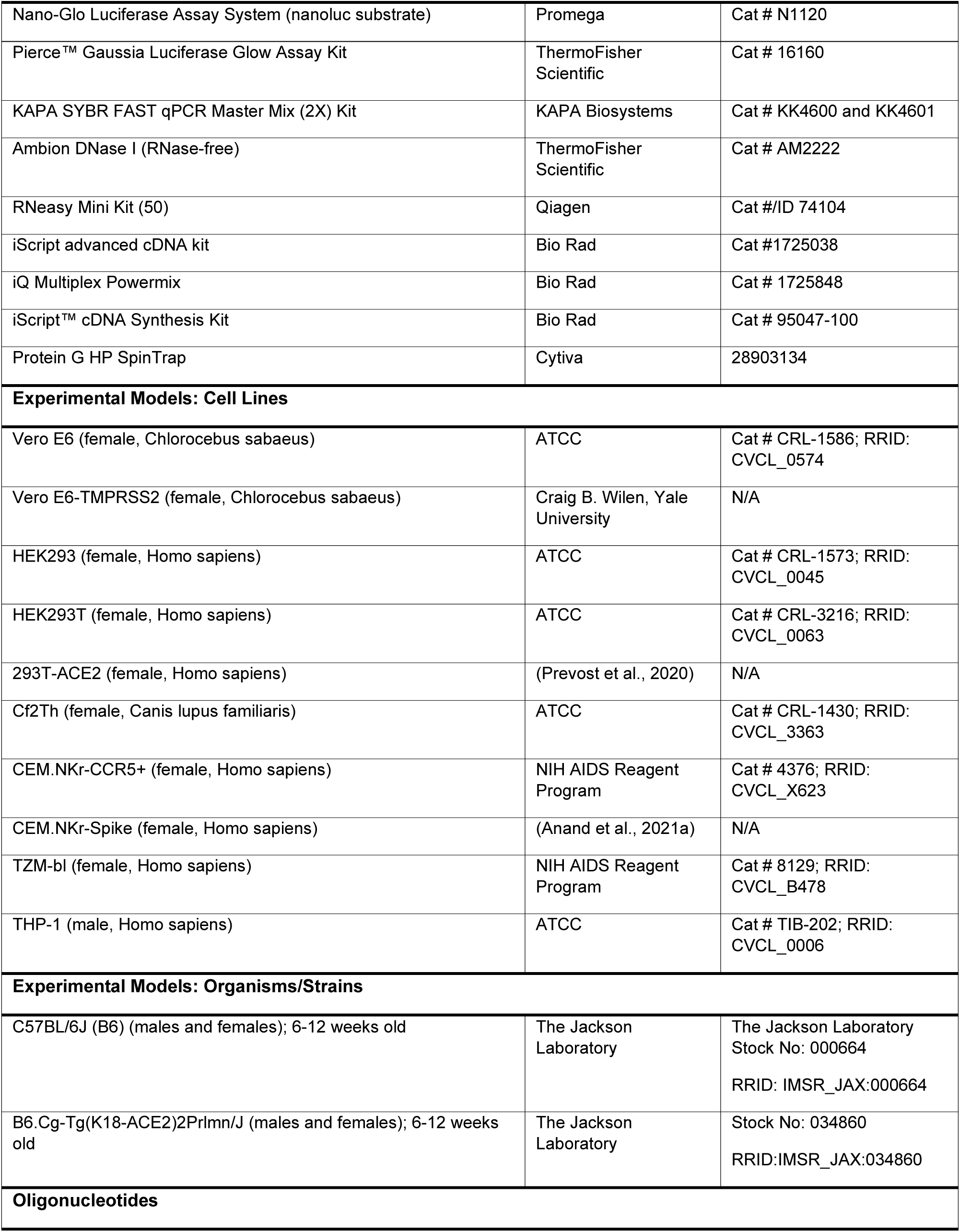
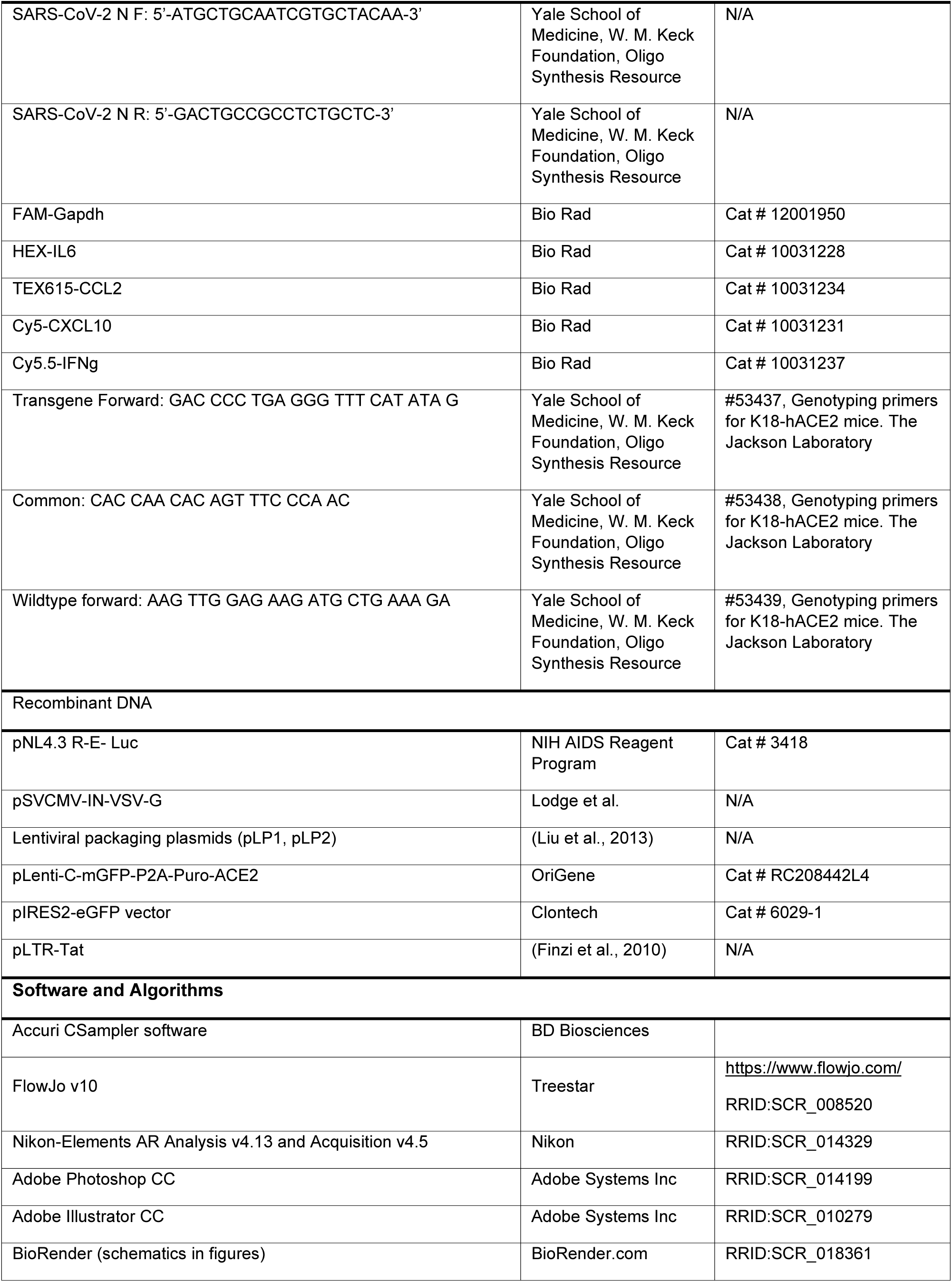
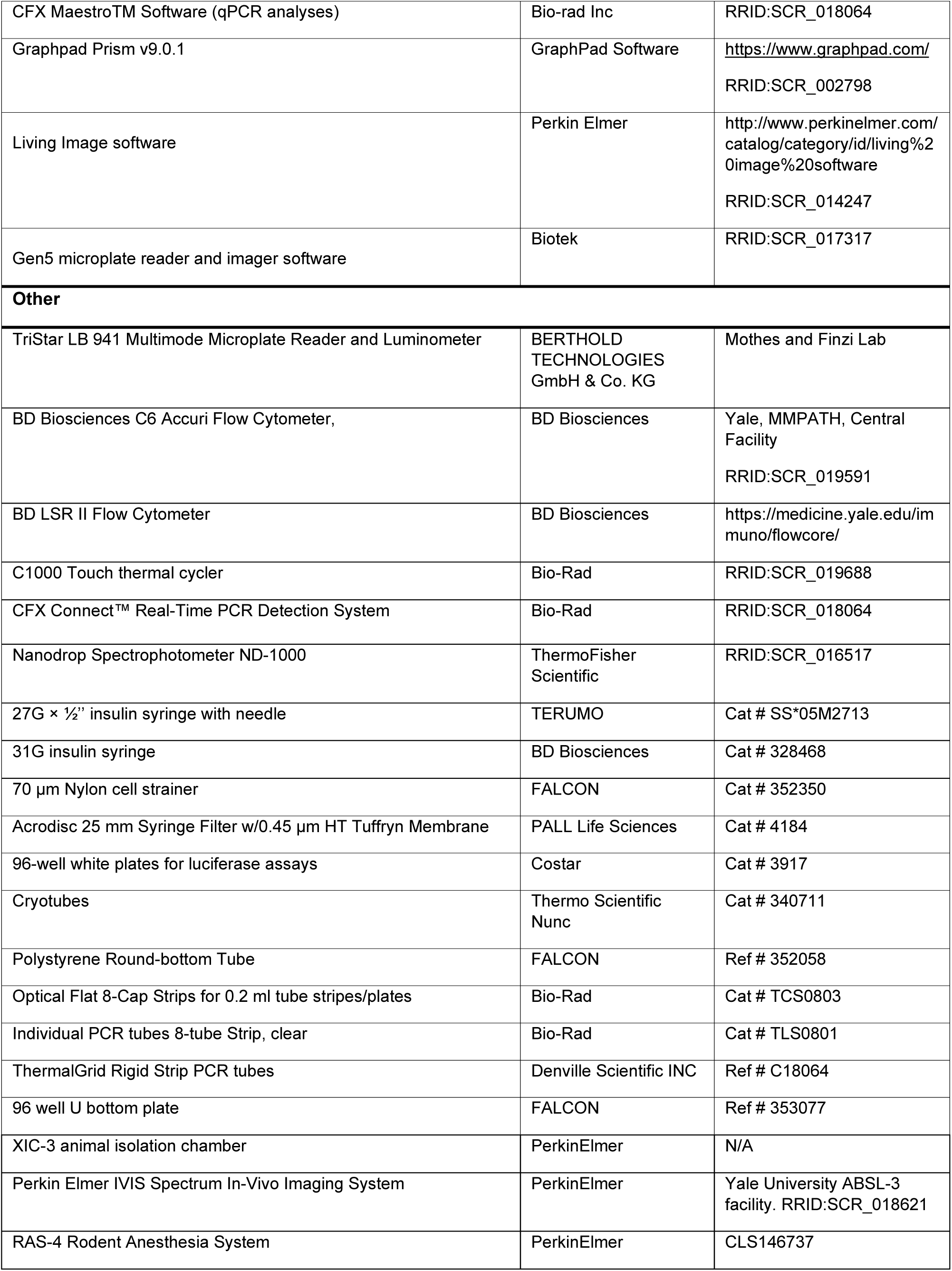
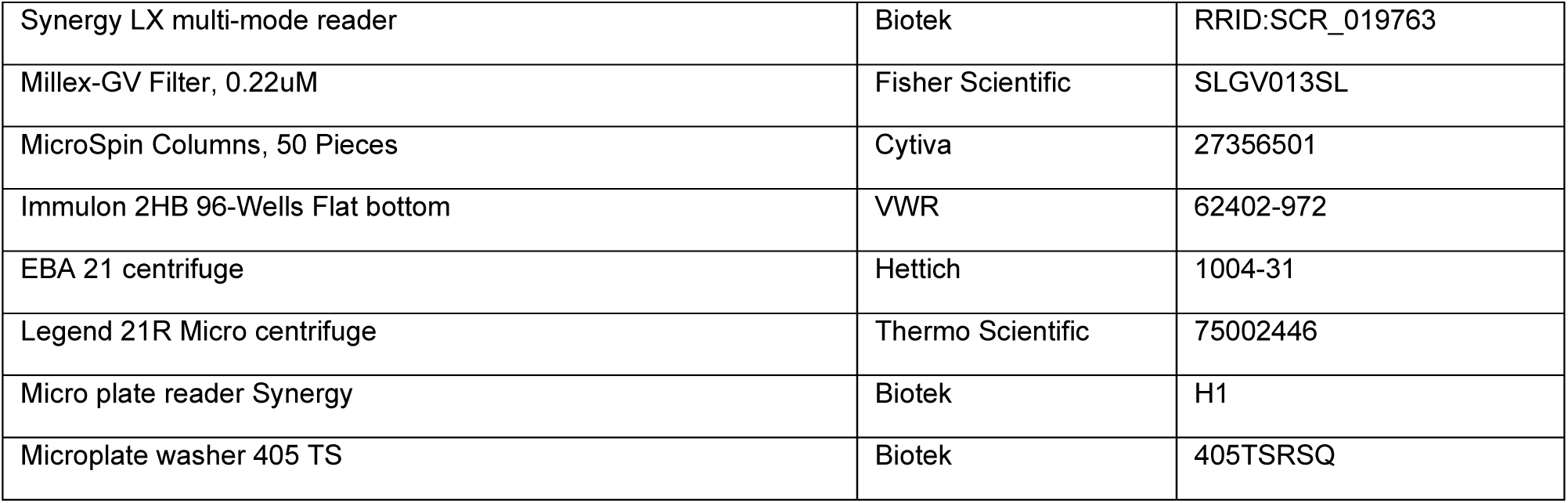

### RESOURCE AVAILABILITY

**Lead Contact:** Pradeep Uchil (

Requests for resources and reagents should be directed to and will be fulfilled by the Lead Contact, Pradeep Uchil,

#### Materials Availability

All other unique reagents generated in this study are available from the corresponding authors with a completed Materials Transfer Agreement.

#### Data and Code Availability

All the data that support the findings of this study are available from the corresponding authors upon reasonable request.

### EXPERIMENTAL MODEL AND SUBJECT DETAILS

#### Cell and Viruses

Vero E6 (CRL-1586, American Type Culture Collection (ATCC), were cultured at 37°C in RPMI supplemented with 10% fetal bovine serum (FBS), 10 mM HEPES pH 7.3, 1 mM sodium pyruvate, 1× non-essential amino acids, and 100 U/ml of penicillin–streptomycin. The 2019n-CoV/USA_WA1/2019 isolate of SARS-CoV-2 expressing Nanoluc luciferase was obtained from Craig B Wilen, Yale University and generously provided by K. Plante and Pei-Yong Shi, World Reference Center for Emerging Viruses and Arboviruses, University of Texas Medical Branch) (Xie et al., 2020). SARS-CoV-2 USA-WA1/2020, B.1.617.2 (Delta) and B.1.351 (Beta) isolates without reporters was obtained through BEI Resources. Viruses were propagated in Vero E6 TMPRSS2 by infecting them in T150 cm^2^ flasks at a MOI of 0.1. The culture supernatants were collected after 72 h when cytopathic effects were clearly visible. The cell debris was removed by centrifugation and filtered through 0.45-micron filter to generate virus stocks. Viruses were concentrated by adding one volume of cold (4 °C) 4x PEG-it Virus Precipitation Solution (40 % (w/v) PEG-8000 and 1.2 M NaCl; System Biosciences) to three volumes of virus-containing supernatant. The solution was mixed by inverting the tubes several times and then incubated at 4°C overnight. The precipitated virus was harvested by centrifugation at 1,500 × g for 60 minutes at 4°C. The concentrated virus was then resuspended in PBS then aliquoted for storage at −80°C. All work with infectious SARS-CoV-2 was performed in Institutional Biosafety Committee approved BSL3 and A-BSL3 facilities at Yale University School of Medicine using appropriate positive pressure air respirators and protective equipment. CEM.NKr, CEM.NKr-Spike, and peripheral blood mononuclear cells (PBMCs) were maintained at 37°C under 5% CO_2_ in RPMI media, supplemented with 10% FBS and 100 U/mL penicillin/ streptomycin. 293T (or HEK293T), 293T-ACE2 (Prevost et al., 2020) cells were maintained at 37°C under 5% CO_2_ in DMEM media, supplemented with 5 % FBS and 100 U/mL penicillin/ streptomycin. CEM.NKr (NIH AIDS Reagent Program) is a T lymphocytic cell line resistant to NK cell-mediated lysis. CEM.NKr-Spike stably expressing SARS-CoV-2 Spike were used as target cells in ADCC assays (Anand et al., 2021; Beaudoin-Bussieres et al., 2021). PBMCs were obtained from healthy donor through leukapheresis and were used as effector cells in ADCC assay.

#### Ethics statement

CCP was obtained from individuals who were infected during the first wave of the pandemic, after at least fourteen days of resolution of COVID-19 symptoms (Perreault et al., 2020). All participants consented to the study (CER #2020-004). PBMCs from healthy individuals as a source of effector cells in our ADCC assay were obtained under CRCHUM institutional review board (protocol #19.381). Research adhered to the standards indicated by the Declaration of Helsinki. All participants were adults and provided informed written consent prior to enrollment in accordance with Institutional Review Board approval.

#### Plasma samples

Recovered COVID-19 patients who have received a COVID-19 diagnosis by the Québec Provincial Health Authority and met the donor selection criteria for plasma donation in use at Héma-Québec were recruited. They were allowed to donate plasma at least 14 days after complete resolution of COVID-19 symptoms. A volume of 500 mL to 750 mL of plasma was collected by plasmapheresis (TRIMA Accel®, Terumo BCT). Disease severity (date of symptoms onset, end of symptoms, type and intensity of symptoms, need for hospitalization/ICU) was documented for each donor using a questionnaire administered at the time of recruitment.

#### Mouse Experiments

All experiments were approved by the Institutional Animal Care and Use Committees (IACUC) of and Institutional Biosafety Committee of Yale University (IBSCYU). All the animals were housed under specific pathogen-free conditions in the facilities provided and supported by Yale Animal Resources Center (YARC). hACE2 transgenic B6 mice (heterozygous) were obtained from Jackson Laboratory. 6–8-week-old male and female mice were used for all the experiments. The heterozygous mice were crossed and genotyped to select heterozygous mice for experiments by using the primer sets recommended by Jackson Laboratory.

### METHOD DETAILS

#### SARS-CoV-2 infection and treatment conditions

For all *in vivo* experiments, the 6 to 8 weeks male and female mice were intranasally challenged with 1 x 10^5^ FFU SARS-CoV-2_WA1_nLuc, WA1, Delta and Beta VOCs in 25-30 µL volume under anesthesia (0.5 - 5 % isoflurane delivered using precision Dräger vaporizer with oxygen flow rate of 1 L/min). For human convalescent plasma treatment using prophylaxis regimen, mice were administered 1 ml of indicated plasma intraperitoneally (i.p.), 24 h prior to infection. For therapy, the same amount was administered two-day post infection (2 dpi). For IgG and Ig(M+A)-depletion, the plasma had to be diluted 1:1. Hence 2 ml of the Class-depleted plasma was administered intraperitoneally in two injections, 1 ml each and 1 hr apart. The starting body weight was set to 100 %. For survival experiments, mice were monitored every 8-12 h starting six days after virus challenge. Lethargic and moribund mice or mice that had lost more than 20 % of their body weight were sacrificed and considered to have succumbed to infection for Kaplan-Meier survival plots. Mice were considered to have recovered if they gained back all the lost weight.

#### IgG and Ig(M+A) depletion of CCP

Selective depletion of IgM, IgA or IgG was done by adsorption on isotype-specific ligands immobilized on sepharose or agarose beads starting with a two-fold dilution of plasma in PBS. IgG and IgA antibodies were depleted from plasma obtained from one recovered COVID-19 patient (CCP6) using Protein G HP Spintrap (GE Healthcare Life Sciences, Buckinghamshire, UK) and Peptide M / Agarose (InvivoGen, San Diego, CA), respectively, according to the manufacturer’s instructions with the exception that no elution step for the recovery of the targeted antibodies was done. For IgM depletion, anti-human IgM (µ-chain specific, Sigma, St.Louis, MO) was covalently coupled to NHS Activated Sepharose® 4 Fast Flow (GE Healthcare) at 815 µg/mL of matrix. Depletion was performed according to the manufacturer’s instructions with the exception that no elution step for the recovery of the targeted isotype was done. All non-depleted and isotype-depleted samples were filtered on a 0.22 µm Millex GV filter (SLGV013SL, Millipore, Burlington, MA) to ensure sterility for the virus capture and neutralization assays. For the preparation of Ig(M+A) depleted samples, plasmas were depleted sequentially in IgM and then in IgA as described above.

To assess the extent of IgM, IgG and IgA depletion, ELISA were performed on non-depleted as well as IgM/IgA- and IgG-depleted plasma samples. Wells of a 96-well microplate were filled with either goat anti-human IgM (µ-chain specific) at 5 µg/mL, goat anti-human serum IgA (a-chain specific) at 0.3 µg/mL or goat anti-human IgG (γ-chain specific) at 5 µg/mL (all from Jackson ImmunoResearch Laboratories, Inc., West Grove, PA). Microtiter plates were sealed and stored overnight at 2-8°C. After four (IgA) to six (IgM and IgG) washes with H_2_O-0.1% Tween 20 (Sigma), 200 μL of blocking solution (10 mmol/L phosphate buffer, pH 7.4, containing 0.85% NaCl, 0.25% Hammerstein casein (EMD Chemicals Inc., Gibbstown, NJ,) were added to each well to block any remaining binding sites. The blocking solution for the IgG and IgM ELISA also contained 0.05% Tween 20. After 0.5 h (IgA) to 1h (IgM and IgG) incubation at 37°C and washes, samples and the standard curves (prepared with human calibrated standard serum, Cedarlane, Burlington, Canada) were added to the plates in triplicates. Plates were incubated for 1h at 37°C. After washes, 100 µL of either goat anti-human IgA+G+M (H+L) HRP conjugate (1/30 000), goat anti-human IgG (H+L) HRP conjugate (1/30 000) or goat anti-human IgA (a-chain specific) HRP conjugate (1/5000) (all from Jackson ImmunoResearch Laboratories, Inc.) were added and samples were incubated at 37°C for 1h. Wells were washed and bound antibodies were detected by the addition of 100 µL of 3,3′,5,5′-tetramethylbenzimidine (TMB, ScyTek Laboratories, Logan, UT). The enzymatic reaction was stopped by the addition of 100 µL 1 N H2SO4 and the absorbance was measured at 450/630 nm within 5 minutes.

#### Bioluminescence Imaging (BLI) of SARS-CoV-2 infection

All standard operating procedures and protocols for IVIS imaging of SARS-CoV-2 infected animals under ABSL-3 conditions were approved by IACUC, IBSCYU and YARC. All the imaging was carried out using IVIS Spectrum® (PerkinElmer) in XIC-3 animal isolation chamber (PerkinElmer) that provided biological isolation of anesthetized mice or individual organs during the imaging procedure. All mice were anesthetized via isoflurane inhalation (3 - 5 % isoflurane, oxygen flow rate of 1.5 L/min) prior and during BLI using the XGI-8 Gas Anesthesia System. Prior to imaging, 100 µL of Nanoluc substrate, furimazine (NanoGlo^TM^, Promega, Madison, WI) diluted 1:40 in endotoxin-free PBS was retroorbitally administered to mice under anesthesia. The mice were then placed into XIC-3 animal isolation chamber (PerkinElmer) pre-saturated with isothesia and oxygen mix. The mice were imaged in both dorsal and ventral position at indicated days post infection. The animals were then imaged again after euthanasia and necropsy by spreading additional 200 µL of substrate on to exposed intact organs. Infected areas identified by carrying out whole-body imaging after necropsy were isolated, washed in PBS to remove residual blood and placed onto a clear plastic plate. Additional droplets of furimazine in PBS (1:40) were added to organs and soaked in substrate for 1-2 min before BLI.

Images were acquired and analyzed with Living Image v4.7.3 *in vivo* software package (Perkin Elmer Inc). Image acquisition exposures were set to auto, with imaging parameter preferences set in order of exposure time, binning, and f/stop, respectively. Images were acquired with luminescent f/stop of 2, photographic f/stop of 8. Binning was set to medium. Comparative images were compiled and batch-processed using the image browser with collective luminescent scales. Photon flux was measured as luminescent radiance (p/sec/cm2/sr). During luminescent threshold selection for image display, luminescent signals were regarded as background when minimum threshold setting resulted in displayed radiance above non-tissue-containing or known uninfected regions. To determine the pattern of virus spread, the image sequences were acquired every day following administration of SARS-CoV-2-nLuc (i.n). Image sequences were assembled and converted to videos using Image J.

#### Focus forming assay

Titers of virus stocks was determined by standard plaque assay. Briefly, the 4 x 10^5^ Vero-E6 cells were seeded on 12-well plate. 24 h later, the cells were infected with 200 µL of serially diluted virus stock. After 1 hour, the cells were overlayed with 1ml of pre-warmed 0.6% Avicel (RC-581 FMC BioPolymer) made in complete RPMI medium. Plaques were resolved at 48 h post infection by fixing in 10 % paraformaldehyde for 15 min followed by staining for 20 min with 0.2 % crystal violet made in 20 % ethanol. Plates were rinsed in water to visualize plaques.

#### Measurement of viral burden

Indicated organs (nasal cavity, brain, lungs) from infected or uninfected mice were collected, weighed, and homogenized in 1 mL of serum free RPMI media containing penicillin-streptomycin and homogenized in 2 mL tube containing 1.5 mm Zirconium beads with BeadBug 6 homogenizer (Benchmark Scientific, TEquipment Inc). Virus titers were measured using three highly correlative methods. Frist, the total RNA was extracted from homogenized tissues using RNeasy plus Mini kit (Qiagen Cat # 74136), reverse transcribed with iScript advanced cDNA kit (Bio-Rad Cat #1725036) followed by a SYBR Green Real-time PCR assay for determining copies of SARS-CoV-2 N gene RNA using primers SARS-CoV-2 N F: 5’-ATGCTGCAATCGTGCTACAA-3’ and SARS-CoV-2 N R: 5’-GACTGCCGCCTCTGCTC-3’.

Second, serially diluted clarified tissue homogenates were used to infect Vero-E6 cell culture monolayer. The titers per gram of tissue were quantified using standard plaque forming assay described above. Third, we used Nanoluc activity as a shorter surrogate for plaque assay. Infected cells were washed with PBS and then lysed using 1X Passive lysis buffer. The lysates transferred into a 96-well solid white plate (Costar Inc) and Nanoluc activity was measured using Tristar multiwell Luminometer (Berthold Technology, Bad Wildbad, Germany) for 2.5 seconds by adding 20 µl of Nano-Glo® substrate in nanoluc assay buffer (Promega Inc, WI, USA). Uninfected monolayer of Vero cells treated identically served as controls to determine basal luciferase activity to obtain normalized relative light units. The data were processed and plotted using GraphPad Prism 8 v8.4.3.

#### Analyses of signature inflammatory cytokines mRNA expression

Brain and lung samples were collected from mice at the time of necropsy. Approximately, 20 mg of tissue was suspended in 500 µL of RLT lysis buffer, and RNA was extracted using RNeasy plus Mini kit (Qiagen Cat # 74136), reverse transcribed with iScript advanced cDNA kit (Bio-Rad Cat #1725036). To determine mRNA copy numbers of signature inflammatory cytokines, multiplex qPCR was conducted using iQ Multiplex Powermix (Bio Rad Cat # 1725848) and PrimePCR Probe Assay mouse primers FAM-GAPDH, HEX-IL6, TEX615-CCL2, Cy5-CXCL10, and Cy5.5-IFNgamma. The reaction plate was analyzed using CFX96 touch real time PCR detection system. Scan mode was set to all channels. The PCR conditions were 95 °C 2 min, 40 cycles of 95 °C for 10 s and 60 °C for 45 s, followed by a melting curve analysis to ensure that each primer pair resulted in amplification of a single PCR product. mRNA copy numbers of *Il6, Ccl2, Cxcl10 and Ifng* in the cDNA samples of infected mice were normalized to *Gapdh* mRNA with the formula ΔC_t_(target gene)=C_t_(target gene)-C_t_(*Gapdh*). The fold increase was determined using 2^-ΔΔCt^ method comparing treated mice to uninfected controls.

#### Antibody depletion of immune cell subsets

Macrophages and neutrophils were depleted during using anti-CSF1R (BioXcell; clone AFS98; 20 mg/kg body weight) (Bauche et al., 2018) and anti-Ly6G (clone: 1A8; 20 mg/kg body weight) (Moynihan et al., 2016) respectively. The mAbs were administered to mice by i.p injection every two days starting at -2 dpi for during CCP prophylaxis or 0 dpi for CCP therapy. Rat IgG2a mAb (BioXCell; clone C1.18.4; 20 mg/kg body weight) or human IgG1 mAb (BioXCell; 12.5 mg/kg body weight) was used as isotype control. The mice were sacrificed and bled 2-3 days after antibody administration or at necropsy to ascertain depletion of desired population.

#### Flow Cytometric Analyses for Immune cell depletion

For analysis of neutrophil depletion, peripheral blood was collected 2-3 days after administration of depleting antibodies. Erythrocytes were lysed with eBioscience 1X RBC lysis buffer (Invitrogen), PBMCs fixed with 4 % PFA and quenched with PBS containing 0.1M glycine. PFA-fixed cells PBMCs were resuspended and blocked in Cell Staining buffer (BioLegend Inc.) containing Fc blocking antibody against CD16/CD32 (BioLegend Inc) before staining with antibodies. Neutrophils were identified as CD45^+^CD11b^+^Ly6G^+^ cells using APC Rat anti-mouse CD45 (30-F11), PE anti-mouse CD11b (M1/70) APC/Cy7 and anti-mouse Ly-6G (1A8) antibodies For analyses of macrophage depletion, lung tissue was harvested 2 days after administration of antibodies. The tissue was minced and incubated in Hanks’ Balanced Salt Solution containing Dispase (5 U/mL; STEMCELL technologies), Liberase TL (0.2 mg/ml, Sigma-Aldrich) and DNase I (100 mg/ml, Roche) at 37°C for 1 h and passed through a 70 µm cell strainer (Falcon, Cat # 352350). The single cell suspension was fixed in BD Cytofix/Cytoperm buffer and stained in BD Cytoperm buffer containing Fc blocking antibody against CD16/CD32 (BioLegend Inc). Macrophages were identified as CD45^+^CD11b^+^Ly6G^-^L6C^-^CD68^+^ population using Alexa 488 Rat anti-mouse CD45 (30-F11), PE anti-mouse CD11b (M1/70) APC/Cy7 and anti-mouse Ly-6G (1A8), APC/Cy7 anti-mouse Ly-6C (HK1.4) and Alexa 647 anti-mouse CD68 (FA-11) antibodies. Data were acquired on an Accuri C6 (BD Biosciences) and were analyzed with Accuri C6 software. 100,000 – 200,000 viable cells were acquired for each sample. FlowJo software (Treestar) was used to generate FACS plot

#### Antibody dependent cellular cytotoxicity (ADCC) assay

This assay was previously described (Anand et al., 2021; Beaudoin-Bussieres et al., 2021). Briefly, for evaluation of anti-SARS-CoV-2 antibody-dependent cellular cytotoxicity (ADCC) activity, parental CEM.NKr CCR5+ cells were mixed at a 1:1 ratio with CEM.NKr cells stably expressing a GFP-tagged full length SARS-CoV-2 Spike (CEM.NKr.SARS-CoV-2.Spike cells). These cells were stained for viability (Aqua fluorescent reactive dye, Invitrogen) and with a cellular dye (cell proliferation dye eFluor670; Thermo Fisher Scientific) and subsequently used as target cells. Overnight rested PBMCs were stained with another cellular marker (cell proliferation dye eFluor450; Thermo Fisher Scientific) and used as effector cells. Stained target and effector cells were mixed at a ratio of 1:10 in 96-well V-bottom plates. Plasma (1/500 dilution) was added to the appropriate wells. Monoclonal antibodies CR3022 and CV3-13 were also included (1 µg/mL) in each experiment as a positive control. The plates were subsequently centrifuged for 1 min at 300 x g, and incubated at 37°C, 5% CO_2_ for 5 hours and then fixed in a 2% PBS-formaldehyde solution. ADCC activity was calculated using the formula: [(% of GFP+ cells in Targets plus Effectors) - (% of GFP+ cells in Targets plus Effectors plus plasma/antibody)]/(% of GFP+ cells in Targets) x 100 by gating on transduced live target cells. All samples were acquired on an LSRII cytometer (BD Biosciences) and data analysis was performed using FlowJo v10.7.1 (Tree Star).

#### Flow cytometry analysis of the different anti-Spike isotypes

For evaluation of the different antibody isotypes (IgG, IgM, IgA and Total Ig) targeting the SARS-CoV-2 Spike, CEM.NKr cells stably expressing a GFP-tagged full length SARS-CoV-2 Spike and CEM.NKr CCR5+ parental cells were stained for 45 minutes at 25°C with plasma CCP-6, plasma CCP-6 depleted in IgG and plasma CCP-6 depleted in IgA and IgM (1/500). Cells were then washed and further stained with a viability dye staining (Aqua fluorescent reactive dye, Invitrogen) and specific secondary antibodies targeting IgGs (Alexa Fluor® 647 anti-human IgG Fc, BioLegend), IgMs (Alexa Fluor® 647-conjugated AffiniPure Goat Anti-Human IgM, Fc5µ Fragment Specific, Jackson ImmunoResearch), IgAs (Alexa Fluor® 647-conjugated AffiniPure Goat Anti-Human Serum IgA, α Chain Specific, Jackson ImmunoResearch) or Total Igs (Alexa Fluor® 647-conjugated AffiniPure Goat Anti-Human IgA + IgG + IgM (H+L), Jackson ImmunoResearch) for 20 minutes at 25°C. The cells were then washed and fixed in a 2% PBS-Formaldehyde solution. The percentage of transduced cells (GFP+ cells) was determined by gating on the living cell population based on the viability dye staining (Aqua fluorescent reactive dye, Invitrogen). Non-specific staining was evaluated using CEM.NKr CCR5+ parental cells and subtracted from the staining on the live GFP+ cells in the CEM.NKr.Spike cells. Samples were acquired on an LSRII cytometer (BD Biosciences) and data analysis was performed using FlowJo v10.7.1 (Tree Star).

#### Pseudovirus neutralization assay

To produce the pseudoviruses, 293T cells were transfected with the lentiviral vector pNL4.3 R-E-Luc (NIH AIDS Reagent Program) and a plasmid encoding for the indicated S glycoprotein (D614G) at a ratio of 10:1. Two days post-transfection, cell supernatants were harvested and stored at -80°C until use. For the neutralization assay, 293T-ACE2 target cells were seeded at a density of 1×104 cells/well in 96-well luminometer-compatible tissue culture plates (Perkin Elmer) 24h before infection. Pseudoviral particles were incubated with several plasma dilutions (1/50; 1/250; 1/1250; 1/6250; 1/31250) for 1h at 37°C and were then added to the target cells followed by incubation for 48h at 37°C. Then, cells were lysed by the addition of 30 µL of passive lysis buffer (Promega) followed by one freeze-thaw cycle. An LB942 TriStar luminometer (Berthold Technologies) was used to measure the luciferase activity of each well after the addition of 100 µL of luciferin buffer (15mM MgSO4, 15mM KPO4 [pH 7.8], 1mM ATP, and 1mM dithiothreitol) and 50 µL of 1mM d-luciferin potassium salt (Prolume). The neutralization half-maximal inhibitory dilution (ID50) represents the plasma dilution to inhibit 50% of the infection of 293T-ACE2 cells by pseudoviruses.

#### SARS-CoV-2 neutralization assay

Serial two-fold dilutions of heat inactivated (56°C for 30 min) CCPs (1:4, 1:16, 1:64, 1:256, 1:1024) were prepared in triplicates in a volume of 50 μL. 50 µL of WA1, Delta and Beta VOCs (a virus concentration to generate 30-50 plaques per well in six well plate) was mixed with diluted plasma and incubated for 1 h at 37°C. The virus-plasma mixes were then added to Vero E6 cells (7.5 × 10^5^ cells/well) seeded 24 h earlier, in 6-well tissue culture plates and allowed to interact with cells for 1 h. The cells were then overlayed with 1 mL of pre-warmed 0.6 % Avicel (RC-581 FMC BioPolymer) made in complete RPMI medium. Plaques were resolved after 72 h by fixing cells in 10 % paraformaldehyde for 15 min followed by staining for 15 minutes with 0.2 % crystal violet made in 20 % ethanol. Plates were rinsed in water to visualize FFU. The FFU counts from virus samples without antibody incubation were set to 100% (30-50 FFU/well). IC_50_ was calculated by plotting the log (plasma dilution) vs normalized FFUs and using non-linear fit option in GraphPad Prism.

#### Quantification and Statistical Analysis

Data were analyzed and plotted using GraphPad Prism software (La Jolla, CA, USA). Statistical significance for pairwise comparisons were derived by applying non-parametric Mann-Whitney test (two-tailed). To obtain statistical significance for survival curves, grouped data were compared by log-rank (Mantel-Cox) test. To obtain statistical significance for grouped data we employed 2-way ANOVA followed by Tukey’s multiple comparison tests. p values lower than 0.05 were considered statistically significant. P values were indicated as ∗, p < 0.05; ∗∗, p < 0.01; ∗∗∗, p < 0.001; ∗∗∗∗, p < 0.0001.

#### Schematics

Schematics for showing experimental design in figures were created with BioRender.com.

## Supplementary Figure Legends

**Figure S1. CCP Efficacy During Prophylaxis and Therapy in K18-hACE2 Mice Against Lethal SARS-CoV-2 Challenge. Related to Figure 2**

(A, B, G, H) Ex vivo imaging of indicated organs and quantification of nLuc signal as flux(photons/sec) after necropsy for an experiment shown in Figure 2A (CCP prophylaxis) and 2G (CCP therapy)

(C, I) Fold change in SARS-CoV-2 nucleocapsid (N gene) expression in brain, lung and nose tissues for experiment as in Figure 2 A and G. The data were normalized to Gapdh mRNA expression in the same sample and that in non-infected mice after necropsy.

(D, J) Viral loads (nLuc activity/mg) from indicated tissue using Vero E6 cells as targets for experiment as in Figure 2 A and G. Undetectable virus amounts were set to 1.

(E, F, K, L) Fold change in cytokine mRNA expression in brain and lung tissues for experiment as in Figure 2 A and G. The data were normalized to Gapdh mRNA expression in the same sample and that in non-infected mice after necropsy.

Viral loads (D, J) and inflammatory cytokine profile (E, F, K, L) were determined after necropsy for mice that succumb to infection at 6 dpi and for surviving mice at 10 dpi (CCP prophylaxis) or 16 dpi (CCP therapy).

Grouped data in (B-F) and (H-L) were analyzed by 2-way ANOVA followed by Tukey’s multiple comparison tests. Statistical significance for group comparisons to isotype are shown in black, with CCP-2 are shown in green, and with CCP-6 are shown in cyan. ∗, p < 0.05; ∗∗, p < 0.01; ∗∗∗, p < 0.001; ∗∗∗∗, p < 0.0001; Mean values ± SD are depicted.

**Figure S2. Immuno-depletion of Neutrophils and Macrophages does not Influence SARS-CoV-2 Replication in K18-hACE2 Mice. Related to Figure 3**

(A, B) Representative FACS plots showing the gating strategy to identify neutrophils (CD45+CD11b+Ly6G+) (n= 8; two experiments) and quantification to ascertain their depletion in blood of indicated groups of mice.

(C, D) Representative FACS plots showing the gating strategy to identify tissue resident macrophages in lungs (CD45+ CD11b+Ly6G-Ly6C-CD68+) (n=8; two experiments) and quantification to ascertain their depletion in single cell suspensions of lung tissue in indicated groups of mice.

(B, D): Non-parametric Mann-Whitney test; ∗, p < 0.05; ∗∗, p < 0.01; ∗∗∗, p < 0.001; ∗∗∗∗, p < 0.0001; Mean values ± SD are depicted.

(E) Experimental design to test effect of macrophages (CD45+Ly6G-Ly6C-CD11b+CD68+) and neutrophils (CD45+CD11b+Ly6G+) depletion in K18-hACE2 mice challenged with SARS-CoV-2- nLuc (1 x 105 FFU). αCSFR-1 or αLy6G mAbs (i.p., 20 mg/kg body weight) were used to deplete macrophages or neutrophils respectively every 48h starting at -2 dpi. Rat isotype mAb treated cohorts served as controls (Iso). Animals were followed by non-invasive BLI every 2 days as indicated.

(F) Representative BLI images of SARS-CoV-2-nLuc-infected mice in ventral (v) and dorsal (d) positions for experiment as in E. Scale bars denote radiance (photons/sec/cm2/steradian).

(G, H) Temporal quantification of nLuc signal as flux (photons/sec) computed non-invasively.

(I) Temporal changes in mouse body weight with initial body weight set to 100% for an experiment shown in E.

(J) Kaplan-Meier survival curves of mice (n = 4 per group; two experiments) for experiment as in E.

**Figure S3. Macrophage and Neutrophil Depletion Compromises CCP-mediated Protection during Therapy Related to Figure 4**

(A-B) Ex vivo imaging of indicated organs and quantification of nLuc signal as flux (photons/sec) after necropsy in K18-hACE2 mice for an experiment shown in Figure 4A

(C) Fold change in SARS-CoV-2 nucleocapsid (N gene) expression in brain, lung and nose tissues. The data were normalized to Gapdh mRNA expression in the same sample and that in non-infected mice after necropsy.

Grouped data in (B-C), were analyzed by 2-way ANOVA followed by Tukey’s multiple comparison tests. Statistical significance for group comparisons to isotype control are shown in black, with CCP-6-treated neutrophil-depleted cohorts are shown in red, CCP-6-treated macrophage-depleted cohorts are shown in green and with CCP-treated cohorts are shown in cyan. ∗, p < 0.05; ∗∗, p < 0.01; ∗∗∗, p < 0.001; ∗∗∗∗, p < 0.0001; Mean values ± SD are depicted. See also Figure 4.

**Figure S4. Characterization of isotype-depleted CCP-6. Related to Figure 5**

(A) Evaluation of indicated Ig class concentration in complete and depleted CCP-6 using ELISA. The relative concentration (%) of each class in the different fractions is shown above the bars.

(B) Levels of Spike-specific denoted antibody classes in complete or depleted CCP-6 fractions, measured by flow cytometry.

(C) Normalized in vitro ADCC activity in complete and depleted CCP-6 fractions evaluated using CEM.NKr cells:CEM.NKr.Spike cells (1:1) as targets and PBMCs from an uninfected donor as effectors.

(D) In vitro neutralization of Spike-decorated lenti-pseudoviral particles by complete and depleted CCP-6 fractions.

(B-D) Undepleted CCP-6 plasma was used for normalizations and set to 100% Mean values ± SD from 3 experiments or replicates is depicted.

**Figure S5 Neutrophils Contribute to Polyclonal IgG-mediated Protection During Prophylaxis in SARS-CoV-2-infected K18-hACE2 Mice. Related to Figure 5**

(A-B) Ex vivo imaging of indicated organs and quantification of nLuc signal as flux(photons/sec) after necropsy for an experiment shown in Figure 5A

(C) Fold change in SARS-CoV-2 nucleocapsid (N gene) expression in brain, lung and nose tissues. The data were normalized to Gapdh mRNA expression in the same sample and that in non-infected mice after necropsy.

Grouped data in (B-C), were analyzed by 2-way ANOVA followed by Tukey’s multiple comparison tests. Statistical significance for group comparisons to isotype control are shown in black, with IgG-equated CCP-6 are shown in cyan, with convalescent plasma IgG depleted CCP-6 are shown in red, with Ig(M+A)-depleted CCP-6 are shown in green and with Ig(M+A)-depleted CCP-6 under neutrophil depletion are shown in orange. ∗, p < 0.05; ∗∗, p < 0.01; ∗∗∗, p < 0.001; ∗∗∗∗, p < 0.0001; Mean values ± SD are depicted.

**Figure S6. Polyclonal IgG and Ig(M+A) Contribute to Protection During CCP Therapy. Related to Figure 6**

(A-B) Ex vivo imaging of indicated organs and quantification of nLuc signal as flux(photons/sec) after necropsy for an experiment shown in Figure 6A

(C) Fold change in SARS-CoV-2 nucleocapsid (N gene) expression in brain, lung and nose tissues. The data were normalized to Gapdh mRNA expression in the same sample and that in non-infected mice after necropsy.

Grouped data in (B-C), were analyzed by 2-way ANOVA followed by Tukey’s multiple comparison tests. Statistical significance for group comparisons to isotype control are shown in black, with IgG-equated CCP-6 are shown in cyan, with IgG-depleted CCP-6 are shown in red and with Ig(M+A)-depleted CCP-6 are shown in green ∗, p < 0.05; ∗∗, p < 0.01; ∗∗∗, p < 0.001; ∗∗∗∗, p < 0.0001; Mean values ± SD are depicted.

