## Supplementary figures and images for "The Fc-effector function of COVID-19 convalescent plasma contributes to SARS-CoV-2 treatment efficacy in mice"

### Supplementary Figures 1-6

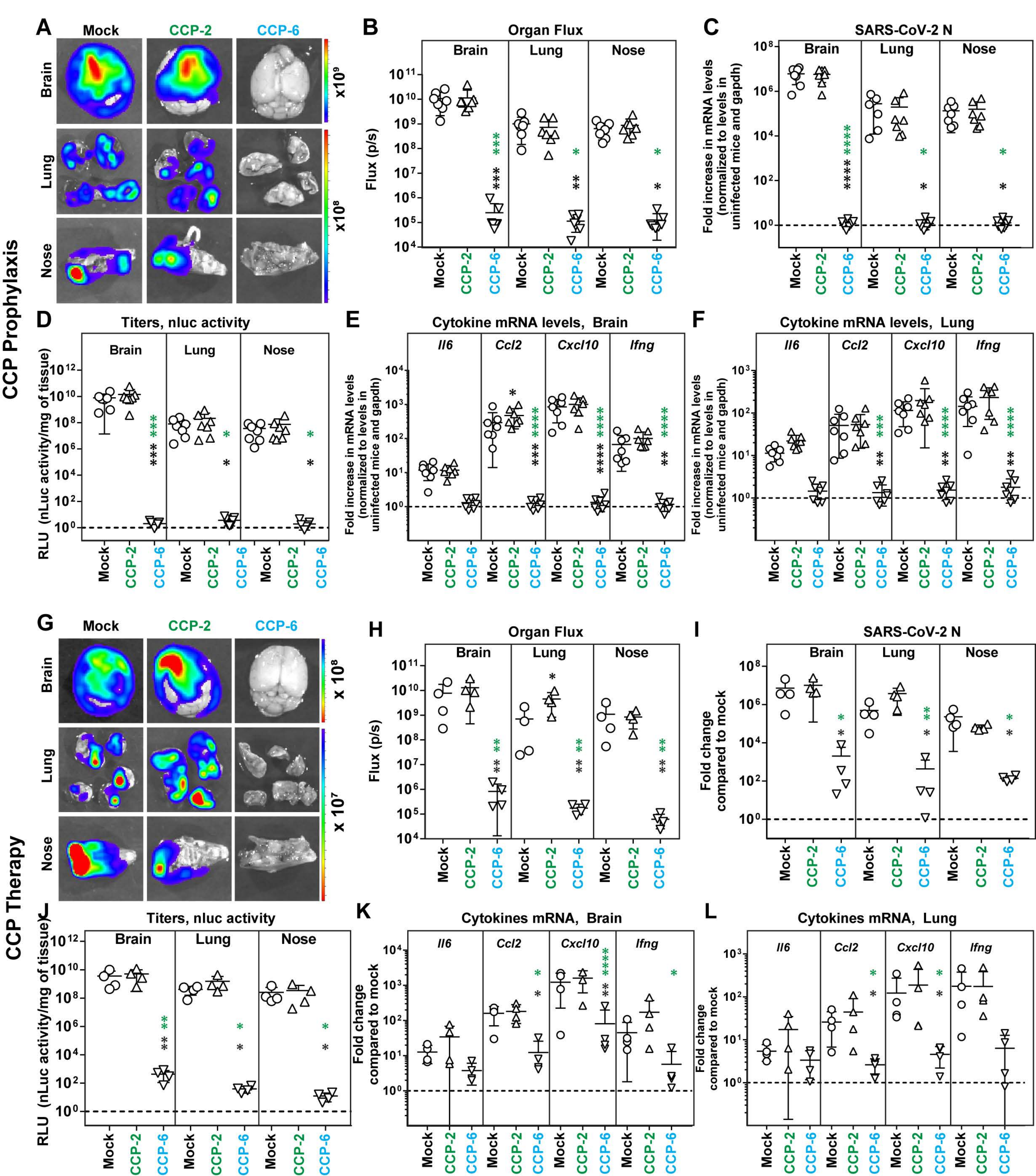

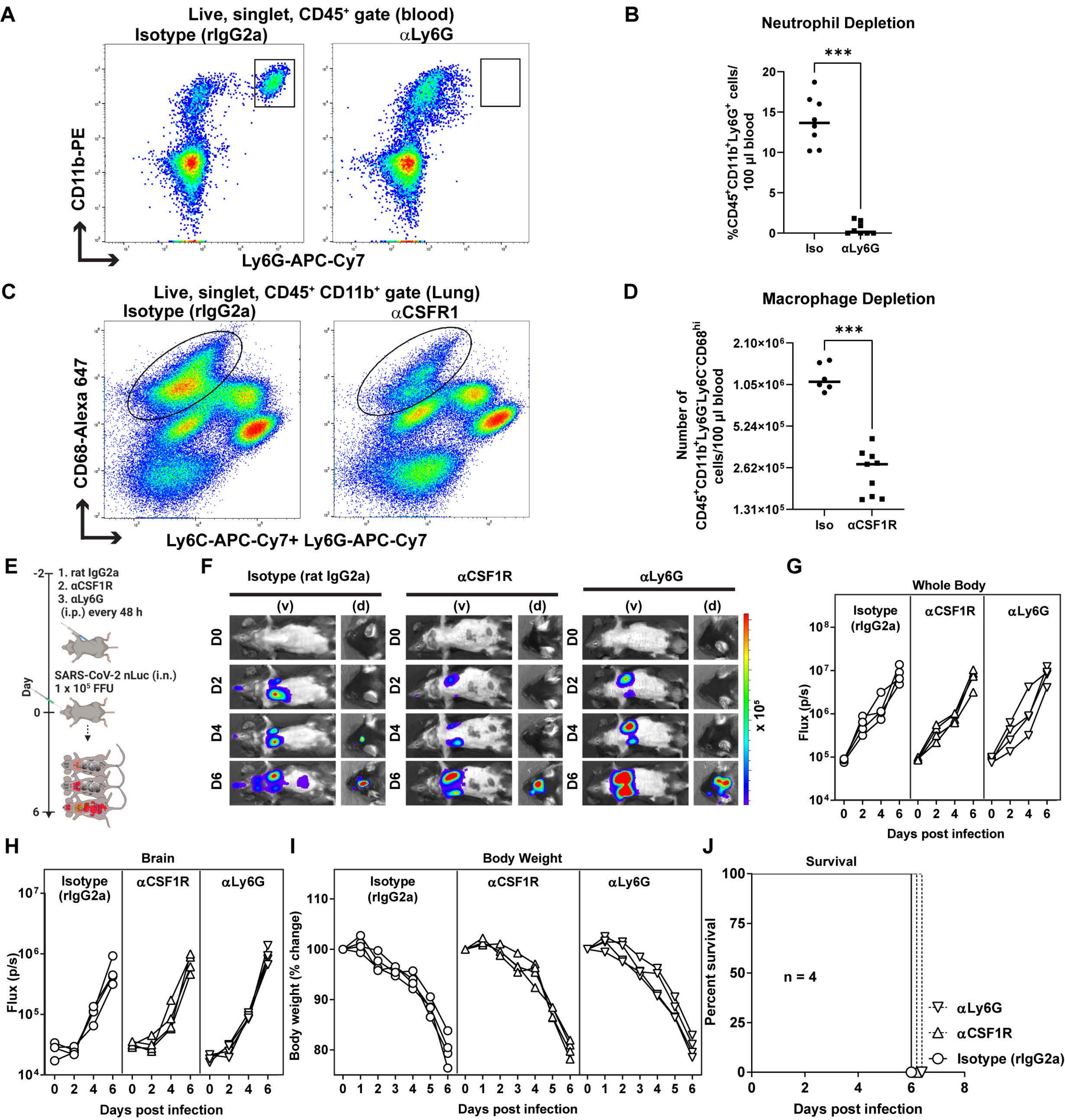

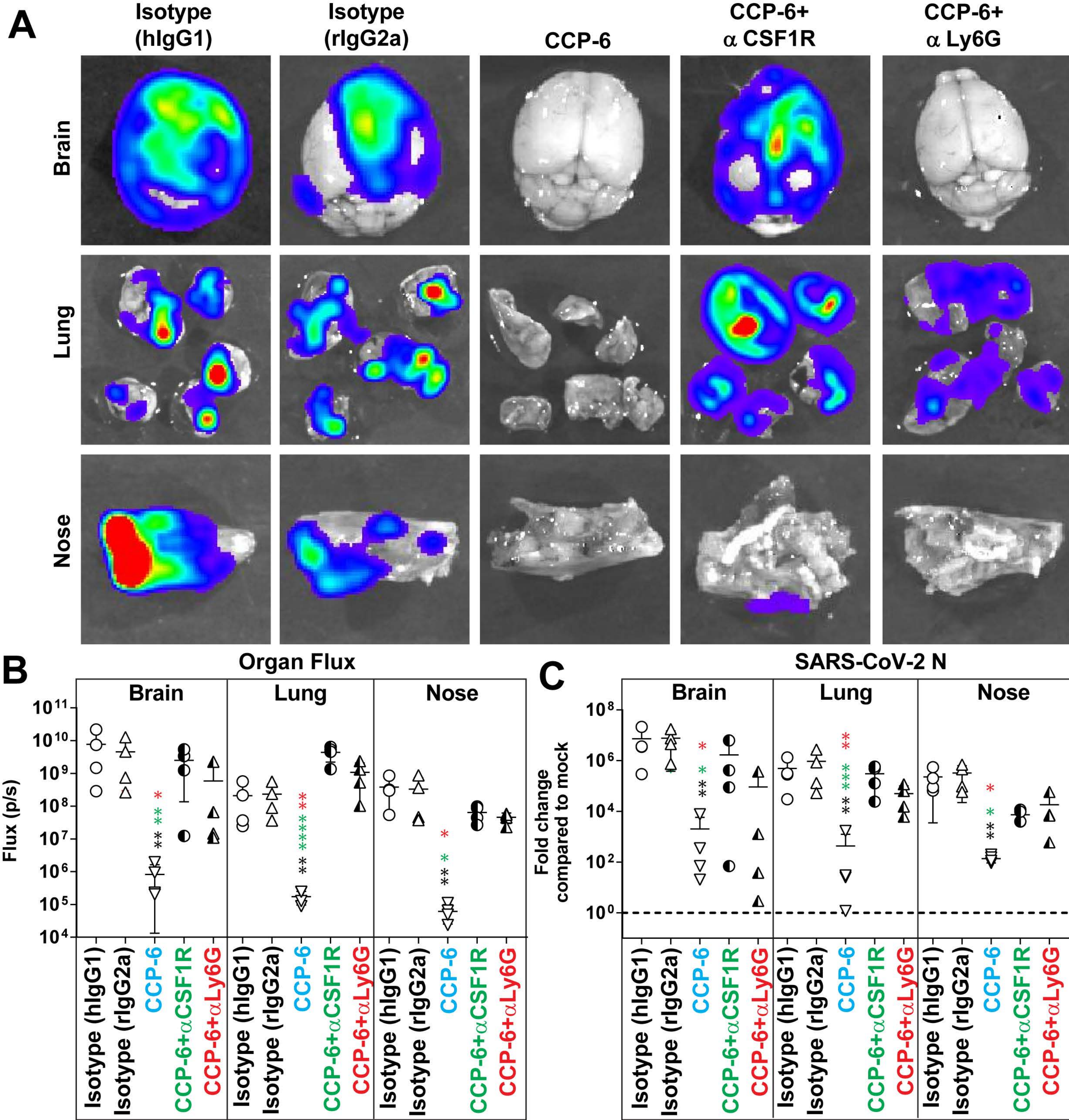

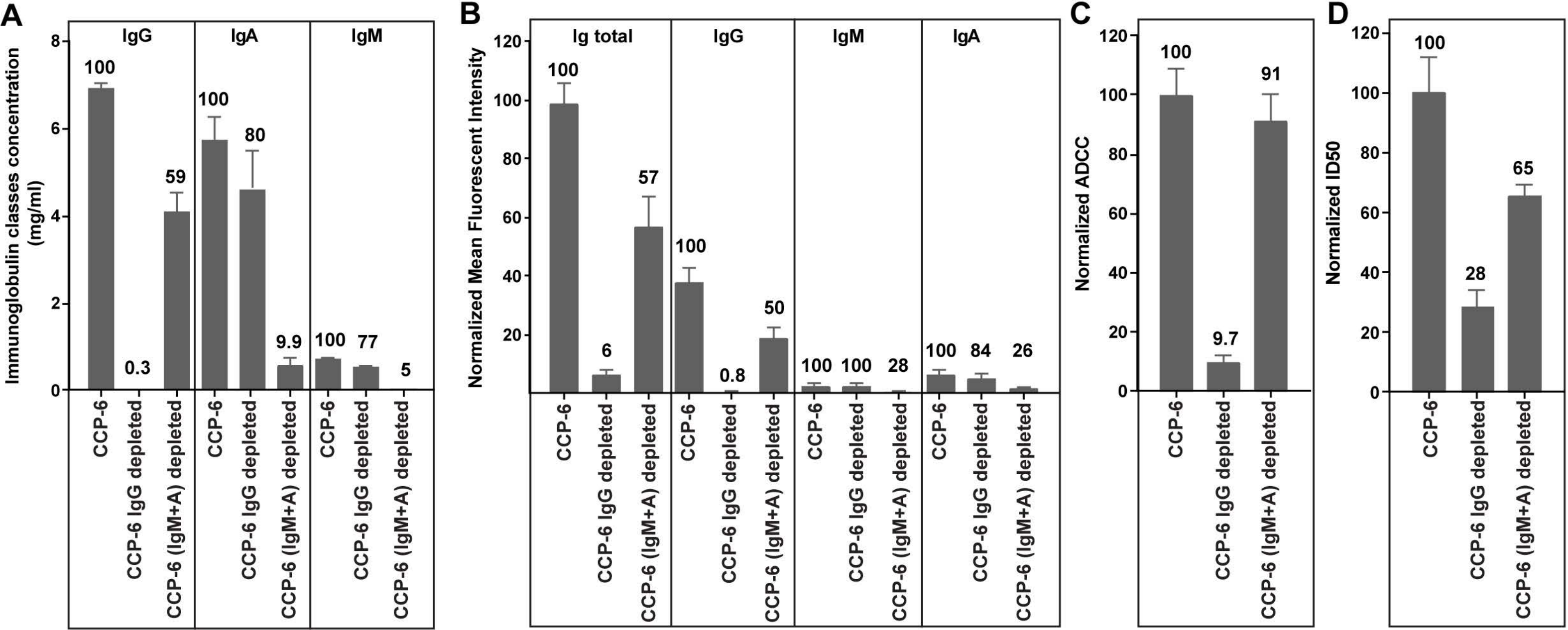

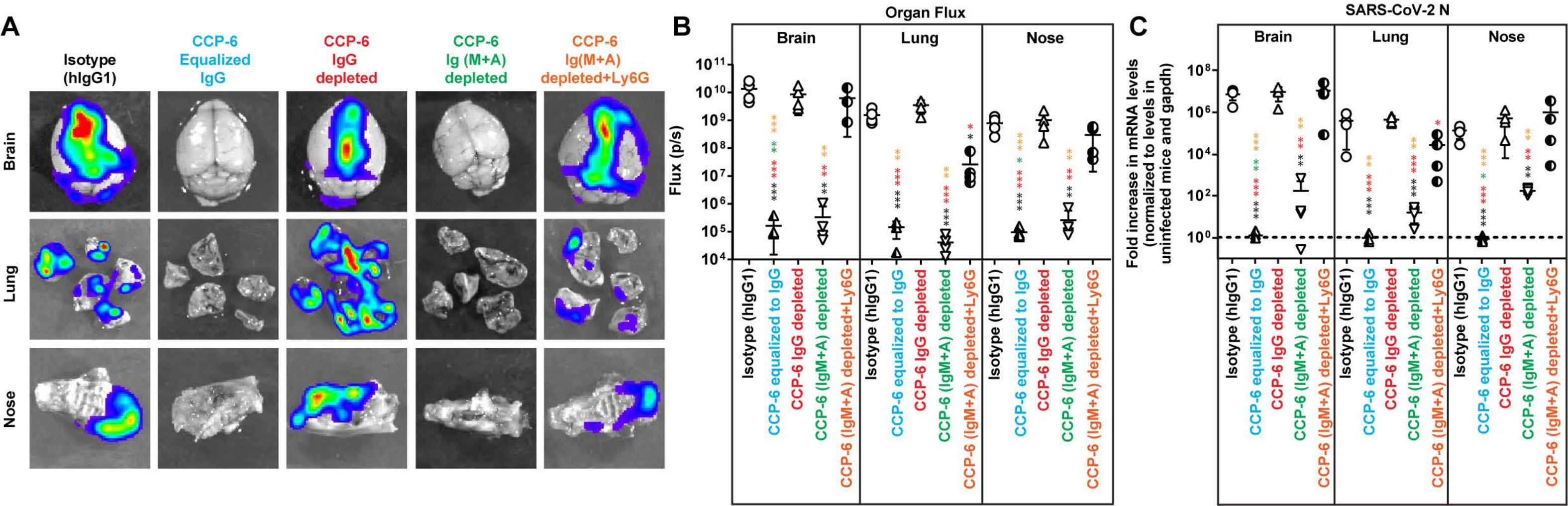

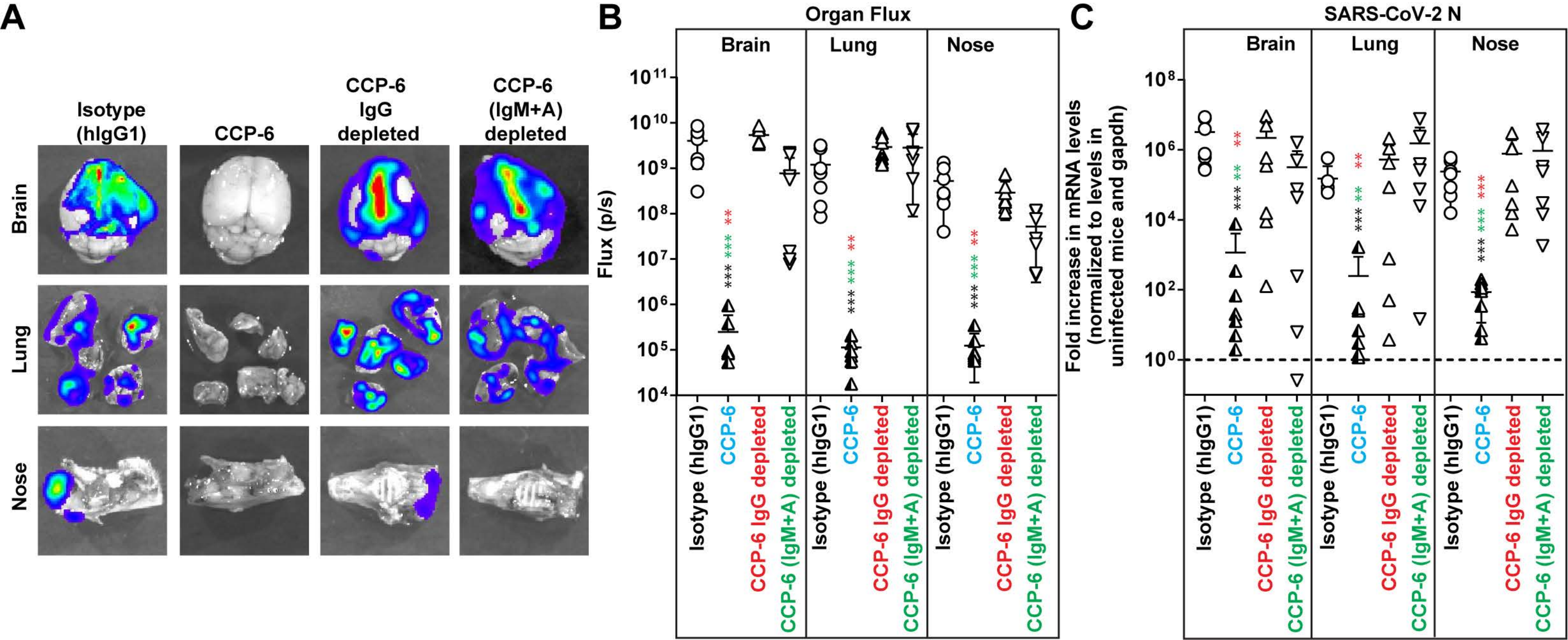
